# Identification, expression, and purification of DNA cytosine 5-methyltransferases with short recognition sequences

**DOI:** 10.1101/2022.06.10.495194

**Authors:** Fumihito Miura, Miki Miura, Yukiko Shibata, Yoshikazu Furuta, Keisuke Miyamura, Yuki Ino, Asmaa M.A. Bayoumi, Utako Oba, Takashi Ito

## Abstract

**Background:** DNA methyltransferases (MTases) are enzymes that induce methylation, one of the representative epigenetic modifications of DNA, and are also useful tools for analyzing epigenomes. However, regarding DNA cytosine 5-methylation, MTases identified so far have drawbacks in that their recognition sequences overlap with those for intrinsic DNA methylation in mammalian cells and/or that the recognition sequence is too long for fine epigenetic mapping. To identify MTases with short recognition sequences that never overlap with the CG dinucleotide, we systematically investigated the 25 candidate enzymes identified using a database search, which showed high similarity to known cytosine 5-MTases recognizing short sequences.

**Results:** We identified MTases with six new recognition sequences, including TCTG, CC, CNG, TCG, GCY, and GGCA. Because the recognition sequence never overlapped with the CG dinucleotide, MTases recognizing the CC dinucleotide were promising.

**Conclusions:** In the current study, we established a procedure for producing active CC-methylating MTases and applied it to nucleosome occupancy and methylome sequencing to prove the usefulness of the enzyme for fine epigenetic mapping. MTases that never overlap with CG dinucleotides would allow us to profile multiple epigenomes simultaneously.

## BACKGROUND

DNA methylation at position 5 of cytosine (5mC) is widely observed in the genomic DNA of living organisms and is one of the major epigenetic modifications. Classically, in mammalian cells, the increased 5mC level was thought to lead to gene silencing, whereas the levels of gene body methylation positively correlate with the expression levels of genes [1–4]. Accordingly, the distribution patterns of 5mC are cell type-specific, and the genome-wide distribution of 5mC or methylome has always attracted much attention.

The deposition of 5mC in mammalian cells depends on three DNA methyltransferases: DNMT1, DNMT3A, and DNMT3B [2, 5, 6]. DNMT1 is a maintenance DNA methyltransferase that introduces methylation to hemimethylated CG dinucleotides. In contrast, DNMT3A and DNMT3B are *de novo* methyltransferases that introduce DNA methylation to non-methylated CG sites. Although some exceptions are known for pluripotent stem cells, oocytes, neurons, and glial cells, DNA methylation on non-CG sites rarely occurs in normal somatic tissues [7, 8]. Therefore, it is commonly accepted that 5mC is applied almost exclusively to CG dinucleotides, and the cytosines in contexts other than CG dinucleotides are unmethylated in most mammalian cells.

The deposition of 5mC is closely related to other epigenetic modifications. For example, 5mC is depleted on the promoters of actively transcribed genes, and trimethylation of lysine 4 of histone H3 (H3K4me3) is enriched [9]. Another example is the di- and tri-methylation of histone H3 lysine 36 (H3K36me2 and H3K36me3), which are positively related to transcriptional activity. H3K36me2 and H3K36me3 are enriched in gene bodies and recruit DNMT3A/B to introduce 5mC. Therefore, these modifications support gene body methylation [10, 11]. Because many epigenetic modifications are mutually related, simultaneous detection of DNA methylation with other epigenetic markers on the same DNA molecule would be useful, and the establishment of such measurement would take the epigenetic measurement to the next level. However, most current methods for detecting epigenomes are unsuitable for simultaneous detection.

For example, chromatin immunoprecipitation followed by sequencing (ChIP-Seq) [12, 13] is the most commonly used technique for detecting epigenomes. In ChIP-Seq, the chromatin is first fragmented, the target epigenome on the fragmented chromatin is then immunoprecipitated, and DNA bound to the epigenome is sequenced. Because ChIP-Seq is based on chromatin fragmentation, the proximally located epigenomes on different nucleosomes should be separated in this first fragmentation step.

Therefore, measuring the relationships between epigenetic modifications on a single DNA molecule is not practical with ChIP-Seq. Accordingly, fragmentation-based procedures are not suitable for measuring multiple epigenetic modifications simultaneously on a single DNA molecule.

There are some fragmentation-free procedures for epigenomic detection. For example, DNA methyltransferase (MTase) can be used as a probe to measure chromatin accessibility. Nucleosome occupancy and methylome sequencing (NOMe-Seq) is one of the first methods to measure chromatin accessibility with an MTase [14], and it has recently been combined with a nanopore sequencer [15]. The same principle was recently applied to Fiber-seq [16], a combination of non-specific adenine N-6 MTase M.EcoGII and a nanopore sequencer. Although chromatin accessibility requires the measurement of DNA-protein interactions of unspecified proteins, interactions between DNA and specific proteins can also be detected with MTase. A representative technology is DamID [17]. In DamID, dam methyltransferase fused with a protein of interest is expressed in target cells, and specific interactions of the fused protein with DNA can be measured by detecting DNA methylation introduced at the N-6 position of adenosine. Although methylation signals were detected using a methylation-sensitive restriction enzyme in the original DamID report [17], this is not the absolute requirement for detecting DNA methylation signals. Indeed, DNA methylation introduced with DamID was detected using single-molecule sequencers. Therefore, MTases enable the fragmentation-free and simultaneous detection of multiple epigenomes on a single DNA molecule.

For epigenetic analysis based on MTases, the resolution is primarily determined by the frequency of the recognized sequences in the genome, and the length of the recognized sequence determines the frequency of sequences. As one of the most basic epigenome units, nucleosomes, are wrapped with approximately 150 base pairs (bp) of genomic DNA, the resolution required for epigenome mapping should be higher than 150 bp. Therefore, the recognition sequence must be 3 bp (64-bp resolution) or less to achieve sufficient resolution for epigenetic measurement. Many MTases have been identified from bacterial and viral sources, and the use of non-specific adenine N6 methyltransferases, such as M. EcoGII, can help achieve fine mapping of chromatin accessibility [16]. However, if we focus on DNA cytosine C-5 MTases, the available MTases that can be used as probes for epigenome mapping are limited to only three in REBASE [18, 19]: M. SssI (<u>C</u>G), M. CviPI (G<u>C</u>), and M. CviPII (<u>C</u>CD) (the target cytosines are underlined). In addition, because of the intrinsic DNA methylation of the CG dinucleotide in mammalian cells, sequences fully (M.SssI) or partially (M.CviPI) overlapping with CG dinucleotides are problematic. Therefore, we launched a project to identify cytosine C-5 MTases with short recognition sequences that never overlap with CG dinucleotides.

The current study presents a systematic strategy for identifying cytosine C-5 MTases in short recognition sequences. This strategy enabled us to identify the MTases of the six novel recognition sequences. In addition, of the identified MTases, we focused on the CC-recognizing MTase because the non-overlapping specificity with CG dinucleotide is beneficial for epigenome mapping of mammalian cells.

## RESULTS

### Identification of cytosine C-5 MTases structurally homologous to M.CviPI and M.CviQIX

We identified two MTases that recognize C-5 cytosine, rather than the CG dinucleotide, in short sequences using REBASE [18, 19] (Figure 1A). The first, M.CviPI (REBASE: 3772, GenBank: AAC64006.1) [20], which could recognize GC dinucleotides, has been commercialized by NEB. The other was M.CviQIX (REBASE: 3175) [21], annotated to methylate the first C of CCD trinucleotide (D is a degenerated expression of A, G, or C).

**Figure 1.**
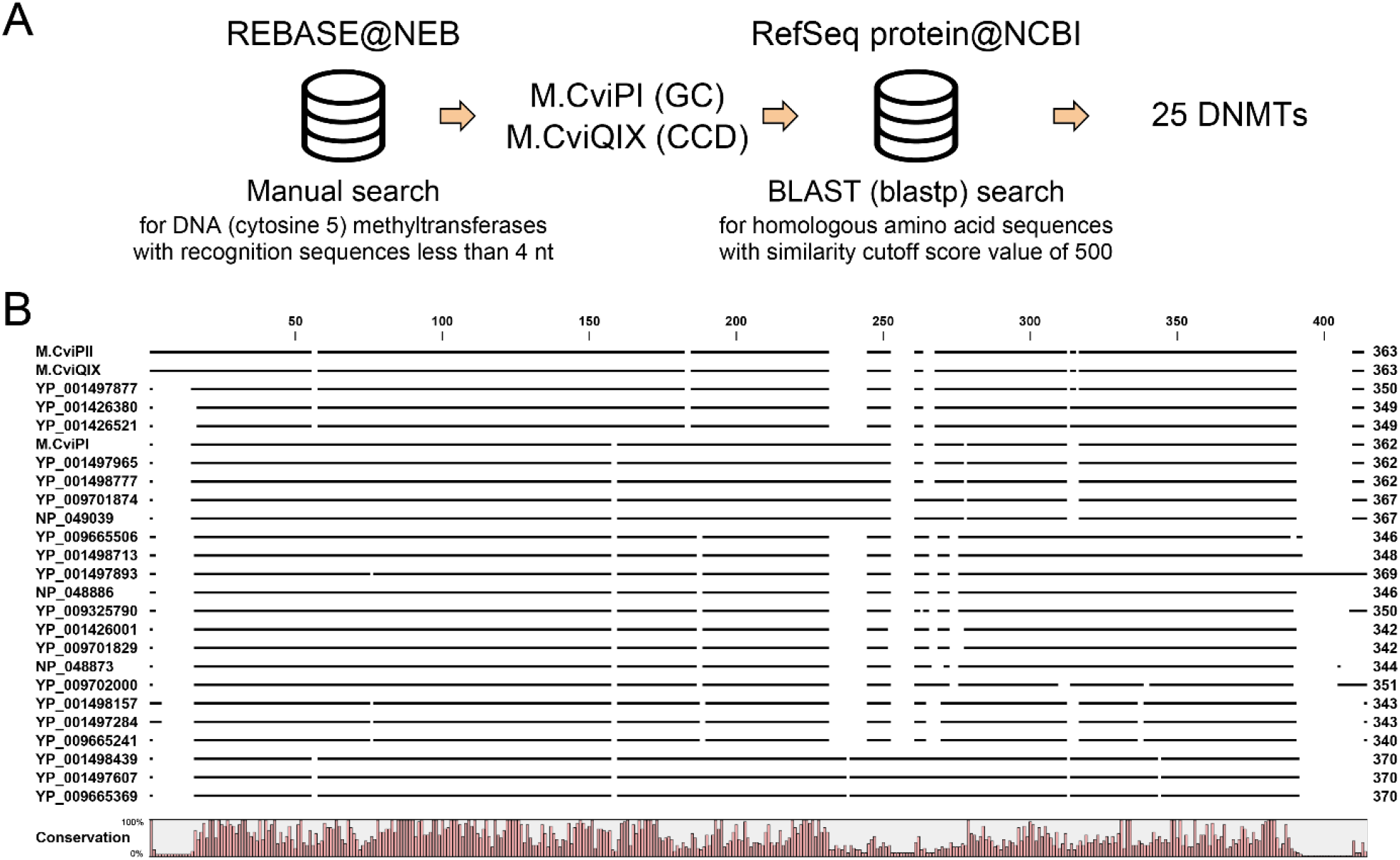
Strategy to identify the MTase genes recognizing short sequence motifs **A**. Scheme used to search the candidate MTases in the databases. **B**. Multiple alignments of the 25 genes. Homologous regions are expressed with thick bars. The same alignment with the single amino acid resolution is provided in Supplementary Figure S2.

In particular, there were some ambiguities in M. CviQIX. The name M.CviQIX was assigned to an unidentified enzyme that might methylate the first C of the CC dinucleotide [21]. Although M.CviQIX was originally isolated from the Chlorella virus strain NY-2A [21], the NY-2A genome sequence (NC_009898) did not contain M.CviQIX. The amino acid sequence of M. CviQIX was unavailable on REBASE, but the sequence we downloaded from REBASE 2 years ago (May 17, 2019) was identical to an MTase found in the genome of a different virus strain AR158 (DQ491003, Gene ID is C619L or RefSeq: YP_001498700). M.CviQIX was almost identical to M.CviPII (GenBank: AAV84097) [22]; only three of 363 amino acids were different between M.CviQIX and M.CviPII. As expected from the high similarity, M. CviPII was identified to methylate the first C of the CCD trinucleotides [22].

M.CviPI and M.CviPII resembled each other [22] but were identified to recognize different short sequences GC and CCD, respectively. Thus, we hypothesized that amino acid sequences structurally homologous to both M. CviPI and M. CviPII (and M. CviQIX) with a similar degree of difference between the proteins might recognize different short sequences. The BLASTP searches [23] using M.CviPI and M.CviPII as queries on non-redundant protein sequences (nr) [24] on the NCBI website identified hundreds of high-scoring hits for both. M.CviPI was identified as a homolog of M.CviPII with a score of 844 and vice versa. Thus, we roughly set the similarity cutoff score as 500. Using these criteria, 25 amino acid sequences homologous to the entire structure of the queries were identified (Figure 1B and Supplementary Figure S1), and the results of two blastp searches shared all genes (not shown). These genes were structurally close, as expected from the procedure used for identification (Figure 1B and Supplementary Figure S1).

Several differences were observed in the multiple alignments of the amino acid sequences (Figure 1B). For example, N-terminal amino acid extension was evident in the CCD-recognizing enzymes M.CviPII and M.CviQIX (Figure 1B and Supplementary Figure S1). Amino acids 217 to 229 of the GC-recognizing enzyme M.CviPI seemed to be insertions (Figure 1B and Supplementary Figure S1). Using a phylogenetic tree based on multiple sequence alignments, we recognized a good diversity of the 25 genes; M.CviPI and M.CviQIX were reasonably separated into different clades (see below).

### A dual-affinity strategy for the expression and purification of active MTases

Before expressing all 25 genes, we investigated *Escherichia coli* protein expression systems using M.CviPI as a model because the successful expression of M.CviPI was previously reported by another group [20]. The cold-shock promoter system was used because it is the primary bacterial protein expression system. Some growth defects were observed after subcloning M.CviPI into the expression vector, even when we used the 5mC-tolerant bacterial strains NEB10β and T7Express that lacked *mcrA, mcrBC*, and *mrr*. The expression of recombinant M.CviPI in bacterial cells was good (Supplementary Figure S2A), and the genomic DNA extracted from the cells was resistant to the methylation-sensitive restriction enzyme HaeIII, which recognized 5’
s-GGCC-3’, after the induction of gene expression (Supplementary Figure S2B). The cold-shock promoter-based expression vector possessed an N-terminal tagged His6 tag so the recombinant protein could be purified via immobilized-metal affinity chromatography (IMAC). Because we wanted to use the M.CviPI combined with antibodies in future studies, fusion proteins of M.CviPI with the antibody-interacting domain of protein A, called ZZ [25], were also constructed. We found that M.CviPI fused with the ZZ domain showed higher expression levels in bacterial cells and higher purity after His6 tag-based purification, especially when ZZ was fused at the C-terminal of M.CviPI (Supplementary Figure S2C), which could be partially explained by the protein solubilization effect of the ZZ domain [26]. Although MTase activity was strong enough with only His-tag-based affinity purification (Supplementary Figure S2C), M.CviPI purified using only the IMAC column with Ni Sepharose resin (HisTrap HP column) contained non-negligible impurities (Supplementary Figure S2D). To further purify the recombinant M.CviPI, we used a dual-affinity purification strategy via fusing the target protein with the Strep-II tag at its C-terminus (Figure 2A). Additional affinity purification of M.CviPI with StrepTactin Sepharose (StrepTrap column) produced proteins with over 95% homogeneity (Figure 2B) with strong DNA methylation activities (Figure 2C). Therefore, we used the cold-shock promoter-based bacterial expression system with the N-terminal His6 tag, ZZ, and Strep-II tag at the C-terminus of the target protein in the following experiments (pCZS vector Supplementary Figure S3A).

**Figure 2.**
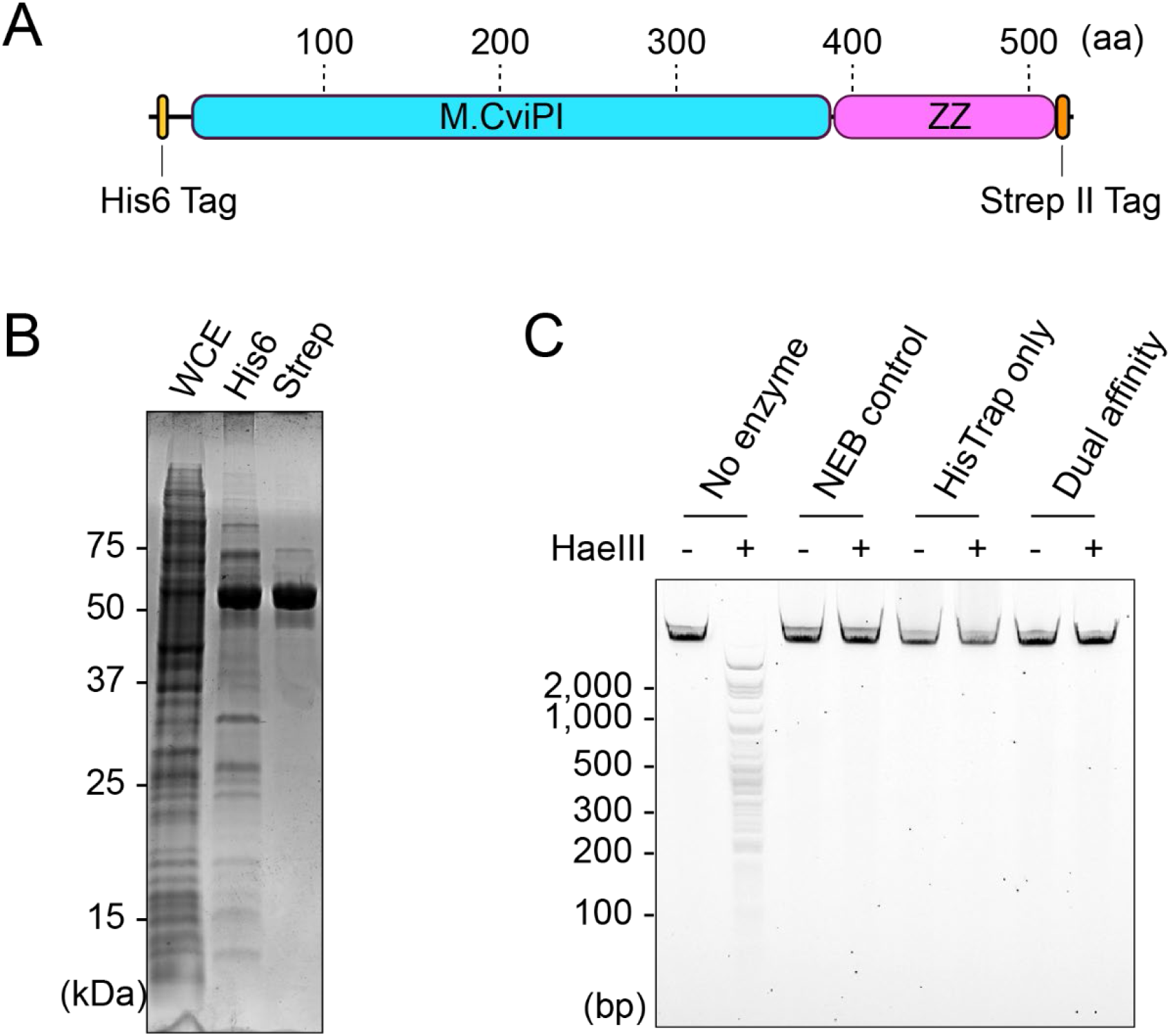
Dual-affinity purification strategy used to purify the candidate MTases. **A**. Structure of the expressed protein. **B**. M.CviPI proteins purified only via HisTrap purification and dual-affinity purification combined with HisTrap and StrepTrap purifications. **C**. DNA methylation activities of purified M.CviPI were assayed using the methylation-sensitive restriction enzyme HaeIII. Unmethylated lambda DNA (Promega) was used as the substrate DNA. As a control, M.CviPI purchased from NEB was included.

### N-terminal deletion enhances the MTase activity of M.CviQIX and M.CviPII

We also tried to produce an active M.CviQIX protein annotated to have MTase activity at the first C of the CCD trinucleotide. If the proteins expressed in the bacterial cells were active, the genomic DNA of the cells would become resistant to both HaeIII and HpaII digestion. Using methylation-sensitive restriction digestion assays, we screened suitable expression systems for M.CviQIX (Figure 3A–C and Supplementary Figure S4A and B). However, we could not detect strong DNA methylation activities in most bacterial cells with genes subcloned into several expression vectors (Figure 3A–C). Strong MTase activity was observed in cells expressing M.CviQIX subcloned in the expression vectors pCTF and pCNS, which produced M.CviQIX fused with a trigger factor (TF) and a SUMO motif at the N-terminal, respectively (Figure 3A). Therefore, we tried to purify the products of these N-terminal TF- and SUMO-tagged constructs. However, both proteins showed only weak MTase activity after affinity purification using the N-terminal His6 tag (not shown).

**Figure 3.**
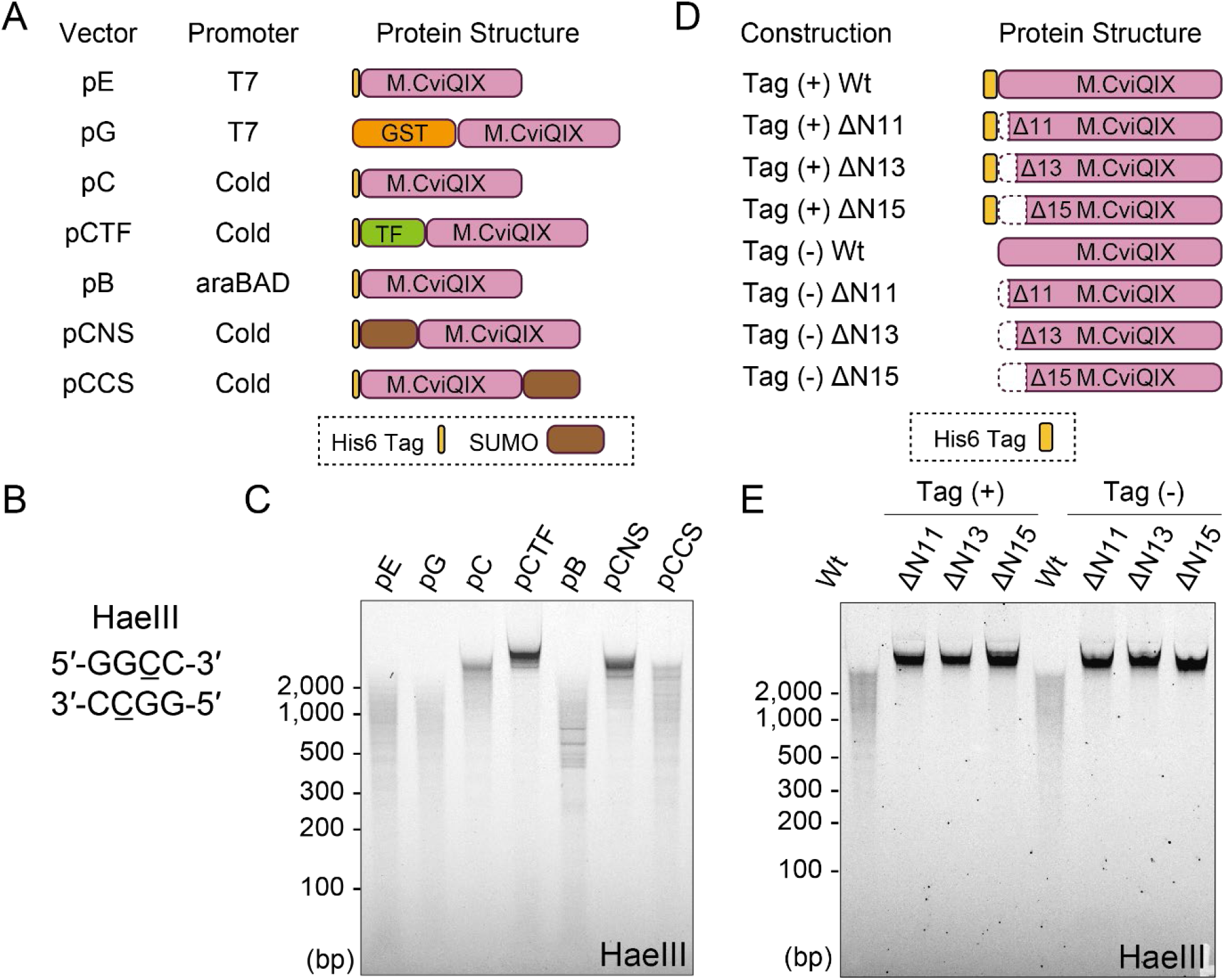
N-terminal extension of M.CviQIX has an inhibitory effect on DNA methylation. **A**. Schemes showing recombinant protein structures of M.CviQIX. **B**. Recognition sequence of the restriction enzyme HaeIII. The recognition sequence contains the CC dinucleotide. If the first C is 5-methylated, HaeIII cannot cut the substrate DNA. **C**. Genomic DNA digestion with HaeIII. DNAs extracted from bacterial cells that express the M.CviQIX proteins shown in A were digested with HaeIII and analyzed on E-Gel Ex. **D**. Schemes showing M.CviQIX proteins with or without N-terminal deletions and vector-derived sequences. Tag denotes the vector-derived sequence. **E**. Genomic DNA digestion of bacterial cells using the constructions in D.

Although SUMO is said to have a protein solubilization effects [27], genomic DNA extracted from the bacterial cells with SUMO-tagged M.CviQIX at the C-terminal showed less resistance to methylation-sensitive restriction enzymes than N-terminally tagged one (Figure 3C), which indicated the N-terminal tagging of the M.CviQIX contributed to the production of active M.CviQIX in the bacterial cells. Additionally, the lack of any MTase activity for the affinity purified N-terminally tagged proteins indicated that the complete N-terminal structure inhibited MTase activity. Furthermore, we observed exceptional amino acid extensions at the N-terminal of both M.CviQIX and M.CviPII (Figure 1B). Therefore, we hypothesized that the N-terminal extensions might cause a low methylation activity of purified M.CviQIX and M.CviPII.

As expected, deletion of the M.CviQIX extension enhanced the methylation of the host genome (Figure 3D and E and Supplementary Figure S4C). Because there were three methionine residues in the N-terminal extension, we speculated that translation initiation might be achieved through one of these residues. Thus, we prepared three N-terminal deletion mutants of M.CviQIX that lacked 11, 13, or 15 amino acids and compared the methylation activities of these proteins (Figure 3D). Genomic DNA extracted from all three deletion mutants showed similar levels of resistance to digestion with the methylation-sensitive restriction enzymes HaeIII and HpaII (Figure 3E and Supplementary Figure S4E). Therefore, any of these three methionine residues could be used as an initiation codon to produce an active M.CviQIX. Note that the N-terminal sequence derived from the expression vector would have a minimal effect on this conclusion because we observed the same result with the constructs possessing no amino acid sequences belonging to the expression vector used (see no tag constructs, Figure 3D and E and Supplementary Figure S4E). We confirmed the same conclusion with the N-terminal deletion mutants of M.CviPII (Supplementary Figure S4F and G). Therefore, the N-terminal amino acid sequences of M.CviQIX and M.CviPII had an inhibitory effect on DNA methylation activity.

### Whole-genome bisulfite sequencing (WGBS) based determination of the MTase recognition sequences

Next, we synthesized the additional 22 MTase genes and aimed to subclone all 25 genes into the expression vector pCZS (Supplementary Figure S3A). We used the 15 amino acid deletion (NDel15) mutants of M.CviQIX and M.CviPII. Fifteen out of the 25 MTase genes were successfully subcloned into pCZS (Supplementary Table S1). In contrast, 10 genes could not be subcloned into the pCZS vector. In addition, we observed that some genes caused severe growth defects after transformation, suggesting leaky expression of the cold-shock promoter of pCZS under uninduced conditions.

Therefore, to enhance the subcloning efficiency, we replaced the cold-shock promoter of pCZS with that of the araBAD system to obtain pBZS (Supplementary Figure S3B) and successfully subcloned all MTase genes into pBZS (Supplementary Table S1).

We induced the expression of these MTases in *E. coli* cells, extracted the bacterial genomic DNA, and performed WGBS using our recent post-bisulfite adapter tagging (PBAT) protocol, tPBAT [28, 29] (Figure 4A). Methylation levels were determined for each cytosine residue. The enrichment of specific nucleobases surrounding the methylated Cs was evaluated by calculating the position weight matrix weighted with methylation levels (PWM-M) and methylation logo (M-logo). For example, as shown in Figure 4B, PWM-M and M-logo for M.CviPI indicated strongly enriched G at the −1 position of C, which indicated that M.CviPI recognized the GC dinucleotide. Using the same strategy, we successfully defined the recognition sequences for 24 out of 25 MTases (Table 1, Supplementary Table S1, and Supplementary Figure S5). Excluding overlaps, these recognition sequences comprised eight unique sequences (T<u>C</u>TG, <u>C</u>G, <u>C</u>C, <u>C</u>NG, T<u>C</u>G, G<u>C</u>Y, G<u>C</u>, and GG<u>C</u>A; the methylated cytosines are underlined). Of these, six (T<u>C</u>TG, <u>C</u>C, <u>C</u>NG, T<u>C</u>G, G<u>C</u>Y, and GG<u>C</u>A) were novel recognition sequences of MTases (Table 1).

**Figure 4.**
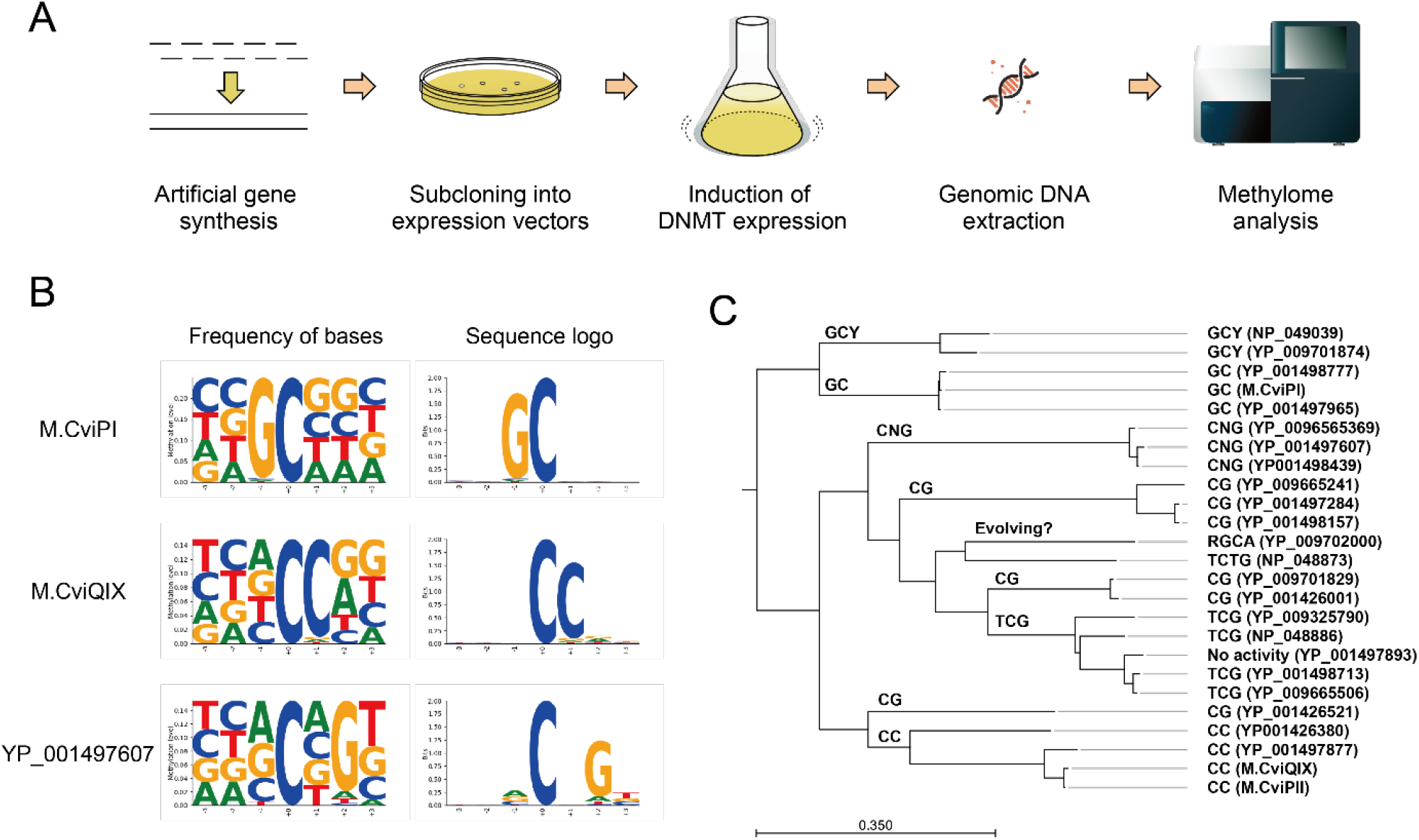
Experimental strategy to define the recognition sequences of the candidate MTase. **A**. Scheme used. **B**. Examples of identified sequence motifs. **C**. Phylogenetic tree was drawn based on the amino acid sequence similarities. The determined recognition sequences were arranged according to the gene name. The common recognition sequence of each clade is annotated at the top of the clade.

**Table 1.**
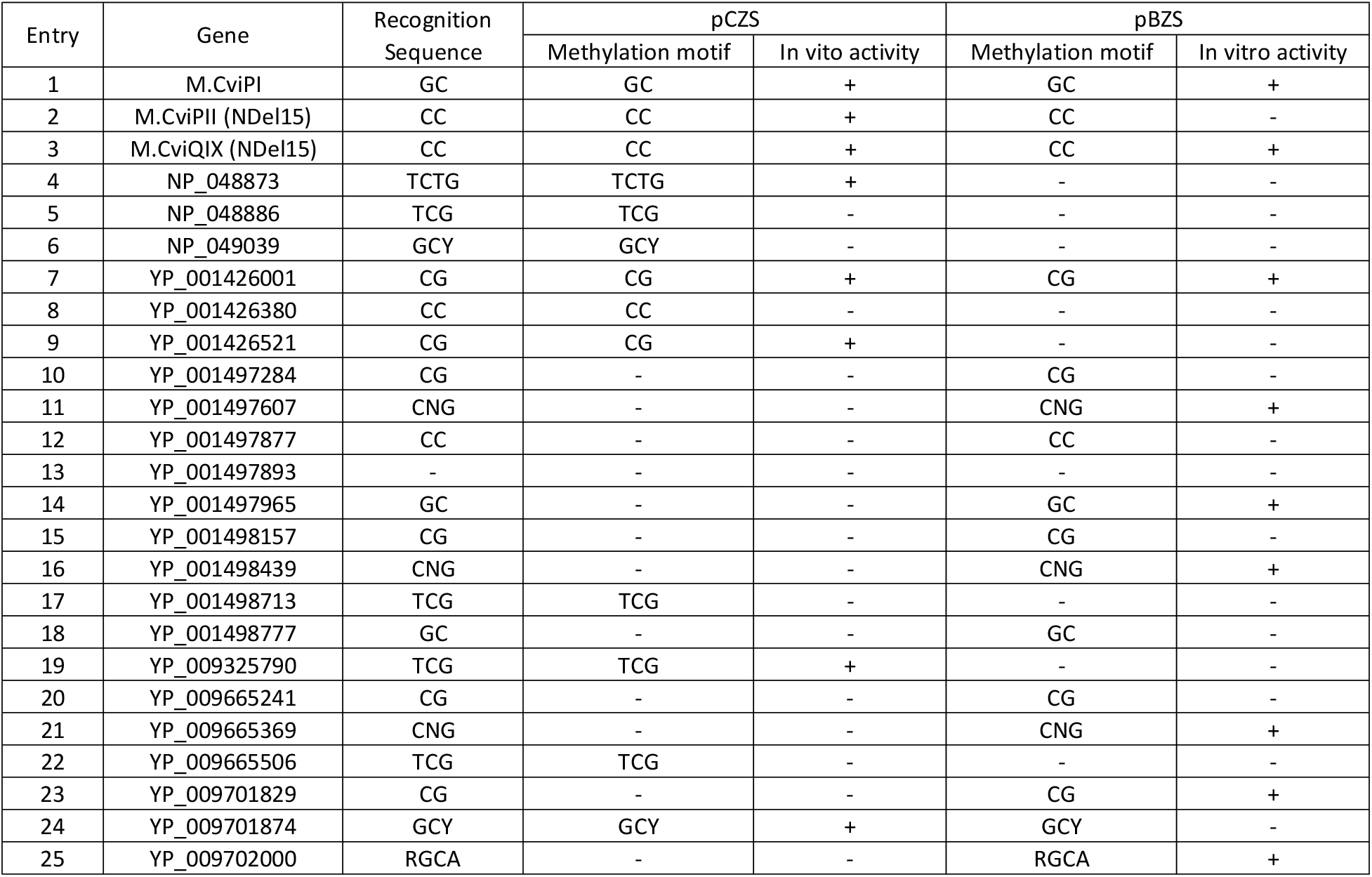
Summary of the analysis of candidate DNMTs.

### Purification of MTases *in vitro*

Next, we attempted to purify the MTases. After cultivating bacterial cells and inducing protein expression, the target MTases were purified using the dual-affinity strategy employing His- and Strep-II-tag (Figure 2). All purifications were performed under the same conditions to systematically evaluate the feasibility of active MTase purification, and the condition used was the one we empirically established to purify the active M.CviPI. Proteins were stored in the presence of 100 µg/mL bovine serum albumin (BSA), 10 mM DTT, and 50% glycerol to preserve enzymatic activity during storage at −20 °C.

Most purified proteins appeared highly homogenous upon SDS-PAGE, as shown in Supplementary Figure S6A. However, it was difficult to obtain sufficient yields of several proteins. We found that some of these MTases showed strong methylation activities when assayed with methylation-sensitive restriction enzymes and/or WGBS using lambda phage DNA as a substrate (Supplementary Table S1 and Supplementary Figure S6B). As a result, 16 active MTases out of 19 were purified (Supplementary Table S1). Some of these proteins lost their activity during purification and storage. We also observed batch-to-batch differences in the activities and purities of these proteins. Therefore, to use these MTases in experiments, further optimization was required for their expression, purification, and storage.

### NOMe-Seq with a CC-recognizing MTase produces similar methylation patterns as GCMT

Of the six newly identified recognition sequences (T<u>C</u>TG, <u>C</u>C, <u>C</u>NG, T<u>C</u>G, G<u>C</u>Y, and GG<u>C</u>A), the CC dinucleotide was promising because the short recognition sequence enabled fine epigenome mapping at approximately 10-bp resolution. Therefore, we aimed to establish an MTase that recognizes CC dinucleotide as a probe for epigenome mapping. We purified the four CC-methylating MTases, M.CviPII, M.CviQIX, YP_001426380, and YP_001497877 (Table 1 and Supplementary Table S1), and found that the N-terminal deletion mutant of M.CviQIX expressed under the control of the cold-shock promoter showed the highest yield and activity (not shown). The purified mutant M.CviQIX is hereafter referred to as CCMT. We investigated the salt concentration in the CCMT reaction and found that 50 mM sodium chloride was the best condition for stringent DNA methylation (Supplementary Figure S7). Notably, M.CviQIX recognizes CC and CCD in the literature [21] and REBASE [18, 19], respectively. Although CCMT recognized every trinucleotide (CCA, CCC, CCG, and CCT) and introduced a methyl moiety to the first C of all (Supplementary Figure S5), we observed slightly weak methylation of CCC trinucleotides in some reaction conditions (Supplementary Figure S7), which might cause a discrepancy between the results reported in the literature and databases.

Next, we used the CCMT for the NOMe-Seq of formaldehyde-crosslinked budding yeast (*S. cerevisiae*) nuclei. Formalin-fixed yeast spheroplasts were permeabilized with an NP-40-containing buffer and treated with CCMT. Genomic DNA was recovered using our recently developed procedure for formalin-fixed samples (Oba U et al., unpublished). Finally, the purified DNA was used for WGBS based on the tPBAT protocol [28]. Because the budding yeast cell did not have any intrinsic DNA methylation, all DNA methylation signals were assumed to be introduced by the MTase treatment. As a control, we performed NOMe-Seq using M.CviPI purchased from NEB (GCMT).

We calculated the methylation levels and compared GCMT- and CCMT-treated nuclei; these two datasets showed similar methylation patterns. (Figure 5A and B, Supplementary Table S2). The aggregation plots of CCMT and GCMT signals centered at several genomic features showed a similar appearance (Figure 5C and D and Supplementary Figure S8A). We observed sharp DNA methylation peaks in the promoter regions of the genes (Figure 5B). These strong DNA methylation signals colocalized well with the signals detected using ATAC-Seq; therefore, these signals were thought to indicate open chromatin regions. The peaks detected upon CCMT- and GCMT-based NOMe-Seq and ATAC-Seq were well-colocalized (Supplementary Figure S8B).

**Figure 5.**
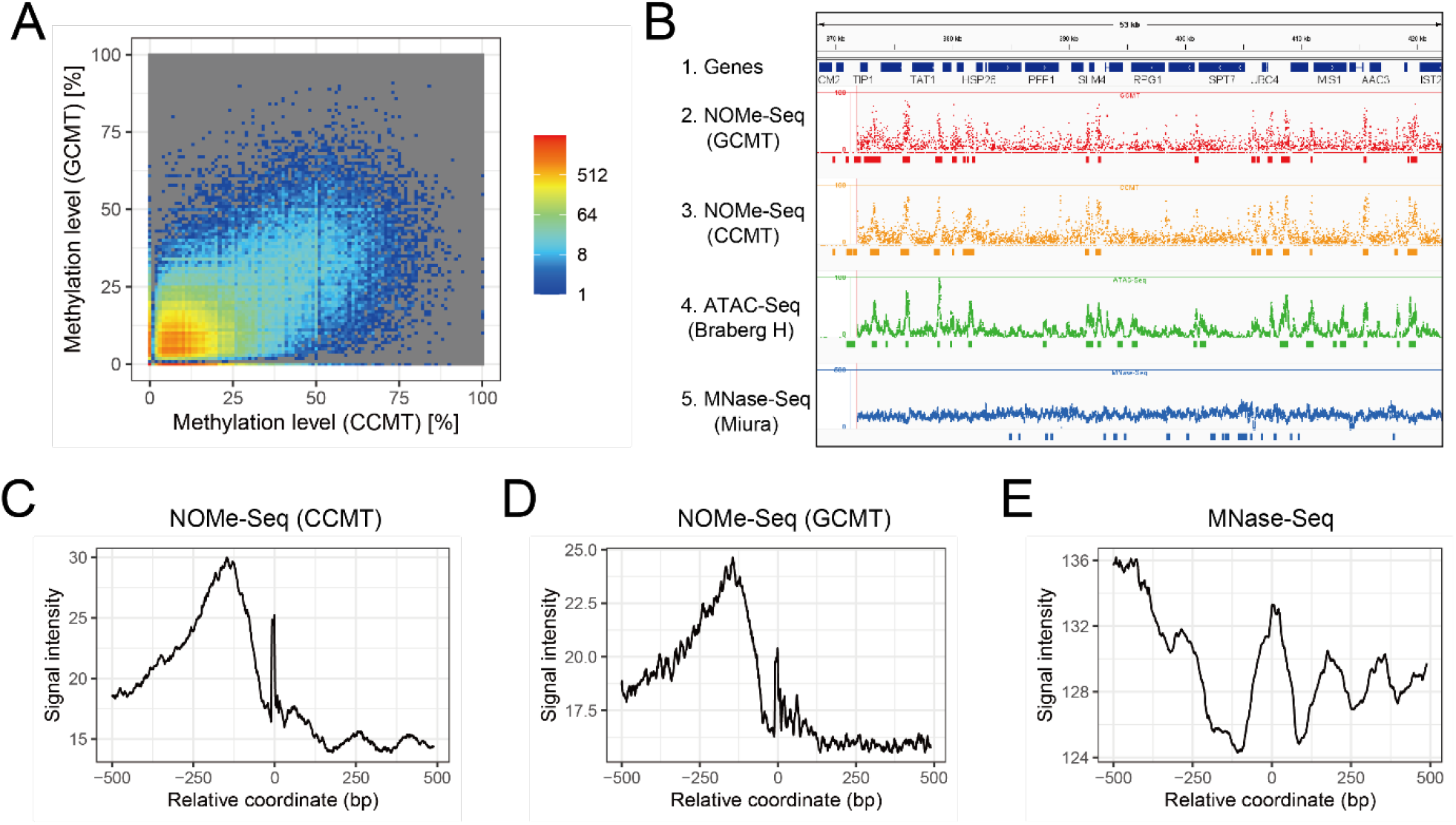
NOMe-Seq of fixed yeast nuclei. **A**. Comparison of methylation levels between CCMT- and GCMT-treated nuclei. Bin size is 10 bp. **B**. Genome browser screenshot showing annotated yeast genes (Track 1), DNA methylation levels at the GC sites (Track 2, NOMe-Seq using GCMT) and CC sites (Track 3, NOMe-Seq using CCMT), read coverage of ATAC-Seq (Track 4, SRR12926697) [36], and MNase-Seq (Track 5, SRR6729489) [37]. **C–E**. Aggregation plots around the initiation codons of yeast genes. DNA methylation signals at CC (C) and GC (D) sites, as well as read coverage of MNase-Seq (E) (SRR6729489), are shown.

The DNA methylation levels at the CC and GC dinucleotides aggregated well at the promoter-proximal regions (Figure 5C and D). Undoubtedly, these signals indicated that nucleosome-free regions were frequently observed at the gene promoters. The promoter-proximal DNA methylation signal at the −100 to −250 region of the transcription start site appeared to be inversely correlated with the low signal region detected via MNase-Seq (Figure 5E). The aggregated DNA methylation signals at the transcription start sites showed a periodicity of 160 to 170 bp, and were inversely correlated with those observed using MNase-Seq (Figure 5B–E). These observations recapitulated the known properties of NOMe-Seq and MNase-seq [30].

### CCMT-based NOMe-Seq achieves fine mapping of the mammalian epigenome

Next, we applied CCMT-based NOMe-Seq to analyze IMR-90 mammalian cells. We obtained high-quality methylome data with 24.5× and 25.7× coverage for CCMT-treated and untreated cells, respectively (Supplementary Table S3). As expected, intrinsic DNA methylation levels at the CG sites were almost identical between the datasets (Figure 6A and B and Supplementary Figure S9A and B). In contrast, DNA methylation levels at the CC sites increased only in the CCMT-treated nuclei (Figure 6C and D). These results indicated that intrinsic CG methylation and CCMT-induced methylation at the CC dinucleotide could be explicitly separated.

**Figure 6.**
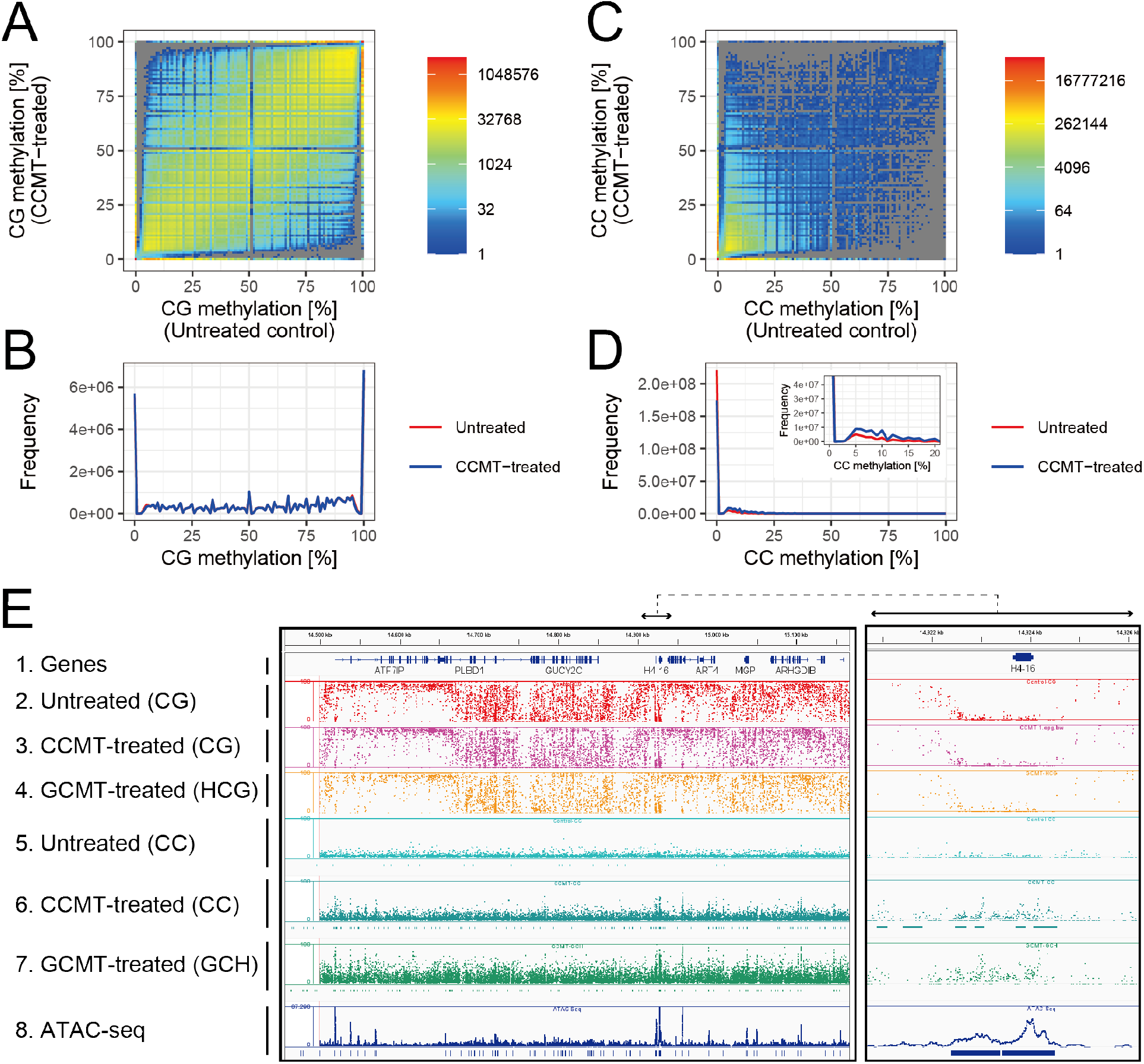
NOMe-Seq of IMR-90 cells. **A–D**. Comparisons of DNA methylation levels between CCMT-treated and untreated nuclei. Intrinsic DNA methylation at the CG sites (A and B) and methylation at CC sites introduced via CCMT *in vitro* (C and D) are shown. **E**. Genome browser screenshots showing annotated human genes (Track 1), intrinsic methylation levels (Track 2, 3, and 4 for untreated, CCMT-treated, and GCMT-treated ones, respectively), MTase-induced methylations (Track 5, 6, and 7 for untreated, CCMT-treated, and GCMT-treated, respectively), and read coverage of ATAC-Seq (SRR7765313) are shown (Track 8). The right panel is the enlarged view of the indicated location of the left panel. For the tracks showing MTase-induced methylation patterns (Track 5, 6, and 7), peaks detected with macs2 are indicated at the bottom of each track.

Then, we visually inspected the two datasets using a genome browser. We observed several large domains with different intrinsic DNA methylation levels at CG sites in both datasets (Figure 6E). These domains were undoubtedly highly methylated regions and partially methylated domains, the existence of which has been described for IMR-90 cells [31]. The comparison between CCMT-based NOMe-Seq and untreated control indicated that these two datasets almost had the same qualities in terms of the intrinsic DNA methylome. In contrast, we observed strong and sharp methylation peaks for CC methylation in the treated samples, whereas none were observed in the untreated control (Figure 6E). These peaks largely colocalized with the peaks identified using ATAC-Seq. Therefore, the peaks observed in the CCMT-induced methylation signals indicated highly accessible chromatin regions (Figure 6E). These results indicated that CCMT-based NOMe-Seq simultaneously measures both the intrinsic methylome and chromatin accessibility of mammalian cells.

We compared the CCMT-based NOMe-Seq with the previously established GCMT-based procedure. We treated IMR-90 cells with GCMT and obtained methylome data with exactly similar procedure conducted for CCMT. We obtained high-quality methylome data with 22.7× coverage (Supplementary Table S3). Note that, to exclude ambiguous GCG sites, only DNA methylation levels in HCG and GCH residues were examined for intrinsic (CG) and artificially induced (GC) DNA methylation, as described by Kelly et al. [14]. The patterns of intrinsic DNA methylation levels were indistinguishable between CCMT- and GCMT-based NOMe-Seq (Figure 6E). In addition, the artificially induced DNA methylation levels at CC and GC sites showed almost similar patterns between the datasets (Figure 6E). Although the methylation levels were higher for GCMT-based NOMe-Seq than that for CCMT-based NOMe-Seq, the methylation levels of NOMe-Seq could be adjusted by fine tuning of the MTase amount used, as previously indicated (15). These results indicated that both CCMT- and GCMT-based NOMe-Seq produce almost similar qualities of data.

To determine whether CCMT-based NOMe-Seq could also detect characteristic aggregation patterns of GCMT-induced DNA methylation around several genomic features [14], we performed the same analysis using the CCMT-based NOMe-Seq data. Methylation patterns around CTCF binding sites were observed in both CCMT- and GCMT-based NOMe-Seq but not in control MTase-untreated WGBS (Supplementary Figure S9C–E). We also found a perk signal in both aggregation plots for GCMT and CCMT around the transcription start sites (Supplementary Figure S9F–H) and histone H3 lysine 4 trimethylation peaks (Supplementary Figure S9I–K). These results also supported the usefulness of CCMT in epigenome mapping.

Ideally, while 233,643,193 GCH sites were available for GCMT-based NOMe-Seq, 302,651,847 sites were usable for epigenome mapping of the human genome with CCMT (Table 2). When Cs covered with at least 10 reads were included in the analysis, 215,834,149 sites were available for CCMT in the WGBS data (see untreated control, Table 2), whereas 165,985,668 sites could be used for GCMT (Table 2). These results indicated that about 30% more sites were available with CCMT than GCMT. The difference in the number of data points directly affected the resolution; the theoretical resolution of the CCMT-based NOMe-Seq (10 bp) was better than that of the GCMT-based NOMe-Seq (13 bp) (Table 2). Although the differences in the analytical resolution between the CCMT- and GCMT-based NOMe-Seq seemed to have little effect on the identification of accessible chromatin regions (Figure 6E), higher analytical resolution would be beneficial for the fine mapping of the epigenetic state of chromatin.

**Table 2.** Resolution of the NOMe-Seq analysis.

| Sequence motif | Genome size (hg19) | Number of motifs in hg19 | Analytical resolution (bp) | Number of motifs with ten or more read coverage |  |  |  |  |  |
| --- | --- | --- | --- | --- | --- | --- | --- | --- | --- |
|  |  |  |  | Control (No MTase) |  | NOME-Seq (CCMT) |  | NOME-Seq (GCMT) |  |
| CG | 3,137,161,264 | 57,400,172 | 55 | 37,593,561 | 65.5% | 35,968,639 | 62.7% | 32,224,228 | 56.1% |
| HCG |  | 43,525,732 | 72 | 28,851,898 | 66.3% | 27,661,596 | 63.6% | 24,764,078 | 56.9% |
| GC |  | 247,517,634 | 13 | 174,727,329 | 70.6% | 169,559,103 | 68.5% | 151,895,283 | 61.4% |
| GCH |  | 233,643,193 | 13 | 165,985,668 | 71.0% | 161,252,061 | 69.0% | 144,435,133 | 61.8% |
| CC |  | 302,651,847 | 10 | 215,834,149 | 71.3% | 208,676,370 | 68.9% | 187,023,053 | 61.8% |

## DISCUSSION

Despite the increasing demand for MTases as tools for mapping the epigenome, the available enzymes are limited, especially for cytosine C5 MTases of short recognition sequences. However, genome sequences have been determined for many bacterial and viral strains. Therefore, we hypothesized that MTases of such short recognition sequences already exist in the databases, although they are still experimentally unidentified.

In this study, we conducted a screening to identify the DNA cytosine C5 MTases with short recognition sequences. We devised a strategy that combines database search, artificial gene synthesis, and WGBS. We successfully determined the recognition sequences of most MTases and purified several MTases with DNA methylation activities. Of the MTases identified, we chose one CC-recognizing enzyme and established it as a new probe for epigenome mapping.

With advanced DNA sequencing and artificial gene synthesis technologies, we have been able to systematically evaluate the recognition sequences of candidate MTases. Similar screenings can be easily set up and applied to any amino acid sequence in the database. During the preparation of this manuscript, Ilinsky et al. showed a similar strategy to determine the recognition sequences of MTases. They combined the expression of artificially synthesized genes in *E. coli* cells and sequenced the bacterial genomic DNA with a nanopore sequencer [32]. In the current study, in contrast, we used WGBS because we thought the reliability of 5mC detection with a nanopore sequencer is currently limited, especially for residues other than the CG dinucleotide. These two successful determinations of recognition sequences using modern DNA sequencing technology would accelerate the identification of unique MTases.

To identify MTases of short recognition sequences, we focused on the mutual sequence similarity of the two identified MTases of different short recognition sequences. We selected structurally similar amino acid sequences in the public database and revealed that these proteins recognize and methylate short sequences. Therefore, the similarity-based approach was effective. Although we have focused on the 25 amino acid sequences registered in RefSeq, many amino acid sequences in the database remain to be identified. Therefore, similar screening should be conducted in the future.

In mammalian cells, cytosine in the CG dinucleotide is methylated by endogenous MTases. In contrast, GCMT methylates cytosine in GC. Based on the difference in the recognition sequences, GCMT has been used for the simultaneous detection of endogenous DNA methylation and chromatin accessibility in NOMe-Seq [14]. However, at the GCG trinucleotide, one cannot discriminate the causative MTases for methylation and, because of this, it is difficult to exploit the full resolution of GCMT for epigenome mapping. Therefore, MTases with non-overlapping recognition sequences with CG dinucleotides would be beneficial. Thus, CCMT is promising because its recognition sequence does not overlap with CG.

The current study has shown the benefits of non-CG overlapping MTases in epigenetic mapping for the simultaneous detection of intrinsic DNA methylation. Similarly, additional MTases that never overlap with CG dinucleotides would expand our ability to profile multiple epigenomes simultaneously. Thus, MTases that recognize CA and CT would be important because these two sequences never overlap with CG and CC. Therefore, we are interested in the two MTases NP_048873 and YP_009702000 that recognize T<u>C</u>TG and GG<u>C</u>A, respectively. Additional screening for genes, particularly homologous to these two MTases, could help identify other MTases that recognize CA and CT residues, which cannot be identified using previous methods.

## METHODS

### Sequence analysis

The amino acid sequences of M. CviPI and M. CviQIX were downloaded from REBASE (http://rebase.neb.com/rebase/rebase.html) [18, 19]. Sequence analyses, including the blastp search, the calculation of multiple alignments, and phylogenetic tree drawing, were performed using the CLC Main Workbench (Qiagen, Hilden, Germany).

### Artificial gene synthesis and subcloning into expression vectors

The genes encoding the MTases were synthesized by Eurofin Genomics Inc. (Tokyo, Japan, for M.CviPI, M.CviPII, and NP_048886) and GenScrypt (Tokyo, Japan, for the other genes) with codon optimization for *E. coli* (Supplementary Information). First, the genes were synthesized with no internal BamHI and EcoRI recognition sequences and with sequences at the 5’ and 3’ terminals, respectively. Then, the gene fragments were subcloned into the BamHI-EcoRI site of pCZS and pBZS (Supplementary Figure S1).

### Small-scale induction of MTases and bacterial genomic DNA extraction

The bacterial expression vectors pCZS and pBZS (Supplementary Figure S3) carrying a gene encoding methyltransferase were introduced into T7Express (New England Biolabs, Ipswich, MA, USA). The cells were inoculated into 3 mL of LB medium (1% [w/v] tryptone, 0.5% [w/v] yeast extract, and 0.5% [w/v] sodium chloride) with 100 µg/mL carbenicillin (Nacalai Tesque, Kyoto, Japan) and grown at 37 °C overnight with vigorous shaking. Fifty microliters of each cell suspension was diluted in 3 mL of LB medium with carbenicillin, and the cells were cultivated at 37 °C for 3 h with shaking. For pCZS, the cell suspension was cooled in ice-cold water for 30 min, and 3 µL of 1 M isopropyl β-D-1-thiogalactopyranoside (IPTG, Nacalai Tesque) was added to induce protein expression. For pBZS, 3 µL of 30% (w/v) L-arabinose was added to induce protein expression. Cells were cultivated with vigorous shaking at 16 °C overnight (pCZS) or 37 °C for 3 h (pBZS). The cells were collected via centrifugation at 2500 ×*g* for 15 min and used for DNA purification using the DNeasy Blood & Tissue Kit (Qiagen), following the optional protocol for gram-negative bacteria provided by the manufacturer. The genomic DNA concentration was determined using the Qubit dsDNA BR Assay Kit and Qubit fluorometer (Thermo Fisher Scientific, Waltham, MA, USA).

### Analysis of DNA methylation with restriction enzyme digestion

Genomic DNA (100 ng) was added to a 10-µL reaction containing 1×CutSmart Buffer and an appropriate restriction enzyme shown in Table 1. The reaction mixture was incubated at 37 °C for 1 h, and the enzyme was heat-inactivated at 70 °C for 10 min. After restriction enzyme treatment, DNA was analyzed using an E-Gel Power SNAP system on a 2% E-Gel Ex gel (Thermo Fisher Scientific).

### Purification of recombinant MTases

The bacterial cells transformed with pCZS or pBZS expressing an MTase-encoding gene were inoculated in 30 mL of LB medium with 100 µg/mL carbenicillin and cultured at 37 °C overnight with vigorous shaking. The seed culture was transferred into a 2-L Erlenmeyer flask with baffles containing 1 L of LB medium with carbenicillin and further cultivated at 37 °C for 4 h, shaking. For pCZS, the flask was cooled in ice-cold water for 30 min, and protein expression was induced by adding 230 mg of IPTG. For pBZS, 3 g of L-arabinose was added to induce protein expression. The flask was incubated at 16 °C overnight (pCZS) or 37 °C for 4 h (pBZS) with vigorous shaking. The cells were collected via centrifugation at 2,500 ×*g* for 15 min, and collected cells were frozen at −80 °C until use. The bacterial cells were resuspended in 20 mL of HisTrap buffer A (20 mM sodium phosphate, pH 7.5, 800 mM NaCl, 20 mM imidazole, 1 mM DTT, and 10% glycerol) containing a protease inhibitor cocktail (Nacalai Tesque) and disrupted via sonication. The lysate was cleared via centrifugation at 15,000 ×*g* for 15 min and filtered using a 0.45-µm syringe filter. The first-round affinity purification was performed using a 5-mL HisTrap HP column (Cytiva, Marlborough, MA, USA) with HisTrap buffer A and HisTrap buffer B (20 mM sodium phosphate, pH 7.5, 800 mM NaCl, 200 mM imidazole, 1 mM DTT, and 10% glycerol). The fraction containing the target protein was further purified using 1-mL StrepTrap HP (Cytiva) with StrepTrap buffer A (100 mM Tris-HCl, pH8.0, 150 mM NaCl, 1 mM EDTA, and 1 mM DTT) and StrepTrap buffer B (00 mM Tris-HCl, pH8.0, 150 mM NaCl, 1 mM EDTA, 1 mM DTT, and 2.5 mM d-desthiobiotin). Chromatographic purification was performed using the AKTA Start System (Cytiva). The dual-affinity purified protein was concentrated via ultrafiltration using an Amicon Ultra-4 30 K device (Millipore), supplemented with BSA and glycerol to obtain final concentrations of 100 µg/mL and 50% (v/v), respectively, and stored at −20 °C until use.

### WGBS

WGBS was performed following the PBAT strategy [29] with recent improvements using a highly efficient single-stranded DNA ligation technique [28]. The molar concentration of the sequencing library was determined using the library quantitation kit from Takara Bio Inc. (Shiga, Japan). Small-scale sequencing was performed using Illumina MiSeq with the MiSeq v3 Reagent Kit in the paired-end mode for 2× 75 cycles (for bacterial and budding yeast cells) or with the MiSeq v2 Reagent Kit nano on the paired-end mode for 2× 150 cycles (for lambda DNA). For large-scale sequencing of mammalian cells, paired-end sequencing for 2× 150 cycles using HiSeq X Ten was performed by Macrogen Japan Corp. (Tokyo, Japan). The reads were mapped to a reference genome (J02459 for lambda phage, NC_000913 for bacterial cells, sacCer3 for yeast cells, and hg19 for human cells) with BMap (https://github.com/FumihitoMiura/Project-2). The mapped alignments were summarized using in-house developed pipelines (https://github.com/FumihitoMiura/Project-2).

### Calculation of the PWM-M and M-logo

From the given genome sequence, a set of every subsequence of length *2l+1* with a C nucleobase at the center of the subsequence was taken, and PWM-M elements were calculated using the following equation:

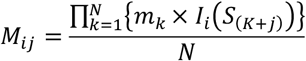

where *i* is nucleobase A, C, G, or T in the matrix, and *j* denotes the relative coordinate from the center of the subsequence. *N* is the total number of C nucleobases in the given genome sequence, and *K* is the genomic coordinate of *k*th C. *S* denotes the nucleobases A, C, G, or T in the genomic sequence, and *S*_*(K+j)*_ indicates the nucleobase at coordinate *K + j. I*_*i*_*(S)* is an indicator function where *I*_*i*_*(S)* is 1 if *S = i*, and 0 otherwise. *m*_*k*_ denotes the methylation level at *k*th C.

From PWM-M, the normalized methylation-weighted frequency *f*_*ij*_ was calculated as follows:

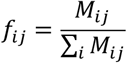

Then, the Shannon entropy *H*_*ij*_ and the information content *R*_*ij*_ of position *j* were calculated, respectively [33, 34]:

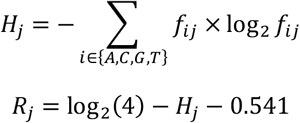

Finally, the height of the logo for each nucleobase *i* at position *j* was calculated as follows:

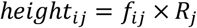

The calculated PWM-M and the M-logo were visualized using the Logomaker [35].

### Budding yeast NOMe-Seq

S288C cells (Biological Resource Center at the National Institute of Technology and Evaluation, NBRC1136) were inoculated into 50 mL of YPD media (1% yeast extract, 2% Bacto peptone, and 2% dextrose) and cultivated with vigorous shaking at 30 °C overnight. Cell suspension was diluted with 1 L of YPD and further cultured at 30 °C for 4 h with shaking. Cells were collected via centrifugation at 2,000 ×*g* for 15 min, washed twice with 50 mL of water, and resuspended in 20 mL of 1 M sorbitol. To the cell suspension, 20 µL of β-mercaptoethanol and 20 mg of Zymolyase 100T (Seikagaku Kogyo Co., Tokyo, Japan) were added and incubated at room temperature for 15 min to degrade the cell wall. Then, the spheroplasts were washed twice with 20 mL of 1 M sorbitol and resuspended in 20 mL of 1 M sorbitol. Fixation was performed by adding 240 µL of 38% formaldehyde (Wako Chemicals, Osaka, Japan) to the resuspended spheroplasts and incubating at room temperature for 5 min. The fixed spheroplasts were washed twice with 20 mL of 1 M sorbitol and pelleted via centrifugation at 5,000 ×*g* for 5 min. The spheroplasts were then resuspended in 10 mL of nuclei preparation solution (10 mM HEPES-KOH, pH7.5, 10 mM NaCl, 3 mM MgCl2, and 0.5% [v/v] Nonidet P 40 substitute) and incubated at room temperature for 5 min. The nuclei were collected via centrifugation at 5,000 ×*g* for 5 min and washed twice with 10 mL of nuclei wash solution (10 mM HEPES-KOH, pH7.5, 10 mM NaCl and 3 mM MgCl_2_). The nuclei were resuspended in 10 mL of nuclei wash solution, of which 500 µL was dispensed into 20 tubes and stored at −80 °C until use.

A frozen nuclei suspension was thawed and resuspended in 500 µL of 1×methylation buffer (50 mM Tris-HCl, pH 8.5, 50 mM NaCl, 1 mM DTT, 1.6 mM SAM). Then, an appropriate amount of either M.CviPI (NEB) or N-terminal 15 amino acid deletion mutant of M.CviQIX (termed CCMT, see RESULTS section), prepared in our laboratory, was added. After incubation at 37 °C for 1 h, the nuclei were pelleted via centrifugation at 5,000 ×*g* for 1 min and resuspended in 40 µL of 10 mM Tris-HCl, pH 8.0. The nuclei suspension was supplemented with 5 µL of 20 mg/mL proteinase K and 10% (w/v) SDS and incubated at 50 °C for 1 h. The reaction was supplemented with 150 µL of 1 M Tris-HCl, pH 8.0, overlayed with white mineral oil, and incubated at 80 °C for 24 h. The aqueous phase was transferred to a new tube and mixed with 200 µL of Buffer AL and isopropanol. The mixture was loaded onto the DNeasy column by centrifuging at 10,000 ×*g* for 1 min, and the column was washed sequentially with 500 µL of Buffer W1 and Buffer W2. Finally, the purified DNA was eluted with 200 µL of Buffer AE and used for the WGBS analysis using the tPBAT protocol. Note that Buffer AL, Buffer W1, Buffer W2, and Buffer AE are DNeasy Blood & Tissue Kit components.

### NOMe-Seq of mammalian cells

The IMR-90 cells (JCRB cell bank #JCRB9054) were cultivated in four T75 flasks containing 30 mL of Minimum Essential Media (Thermo Fisher Scientific) supplemented with 10% fetal bovine serum (Thermo Fisher Scientific) and 50 U/mL penicillin-streptomycin (Thermo Fisher Scientific) at 37 °C in an atmosphere of 100% humidity and 5% CO_2_. Cells grown were treated with trypsin and collected via centrifugation at 2,000 ×*g* for 5 min. Cells were washed twice with 20 mL of D-PBS (Nacalai Tesque) and resuspended in CELLBANKER 1 (Takara Bio) at 1 × 10^7^ cells/mL. The cell suspension was dispensed in 500 µL and stored at −80 °C until use.

Before use, the frozen cell suspension was thawed and washed twice with PBS. Cells were then resuspended in nuclei preparation buffer (10 mM Tris-HCl, pH 8.0, 10 mM NaCl, 3 mM MgCl2, 0.1 mM EDTA) and 0.5% (w/v) NP-40, and incubated on ice for 10 min. The nuclei were collected via centrifugation at 2,000 ×*g* for 5 min, resuspended in with 500 µL of 1× methylation buffer with an optimized amount of CCMT, and incubated at 37 °C for 30 min. Then, the nuclei were collected via centrifugation at 2,000 ×*g* for 5 min, and the supernatant was removed. The nuclei were dissolved in 20 µL of 20 mg/mL proteinase K and 200 µL of Buffer ATL was added. The reaction was incubated at 56 °C for 1 h and supplemented with 200 µL of Buffer AL and 200 µL of isopropanol. The mixture was loaded onto the DNeasy column and washed with 500 µL of buffer W1 or Buffer W2. The purified DNA was eluted with 200 µL of Buffer AE and used for WGBS.

## LIST OF ABBREVIATIONS

BSA: Bovine serum albumin
IMAC: Immobilized-metal affinity chromatography
PBAT: Post-bisulfite adapter tagging
TF: Trigger factor
WGBS: Whole-genome bisulfite sequencing

## DECLARATIONS

### Ethics approval and consent to participate

Not applicable.

### Consent for publication

Not applicable.

### Availability of data and materials

All sequence data used in this study were deposited to NCBI GEO and NCBI SRA with accession numbers GSE192699, GSE192949, GSE202353, GSE202354, and SRR17348378-SRR17348416.

### Authors’ contributions

FM conceived the study, performed experiments, analyzed data, and wrote the paper. MM, YS, KM, YI, AB performed experiments. YF and UO contributed to experimental design. TI supervised the study.

## Acknowledgments

We would like to thank Editage (www.editage.com) for English language editing.

## SUPPLEMENTARY DATA

Supplementary Data are available online.

