## Supplementary Information for "Identification, expression, and purification of DNA cytosine 5-methyltransferases with short recognition sequences"

### Supplementary Figures

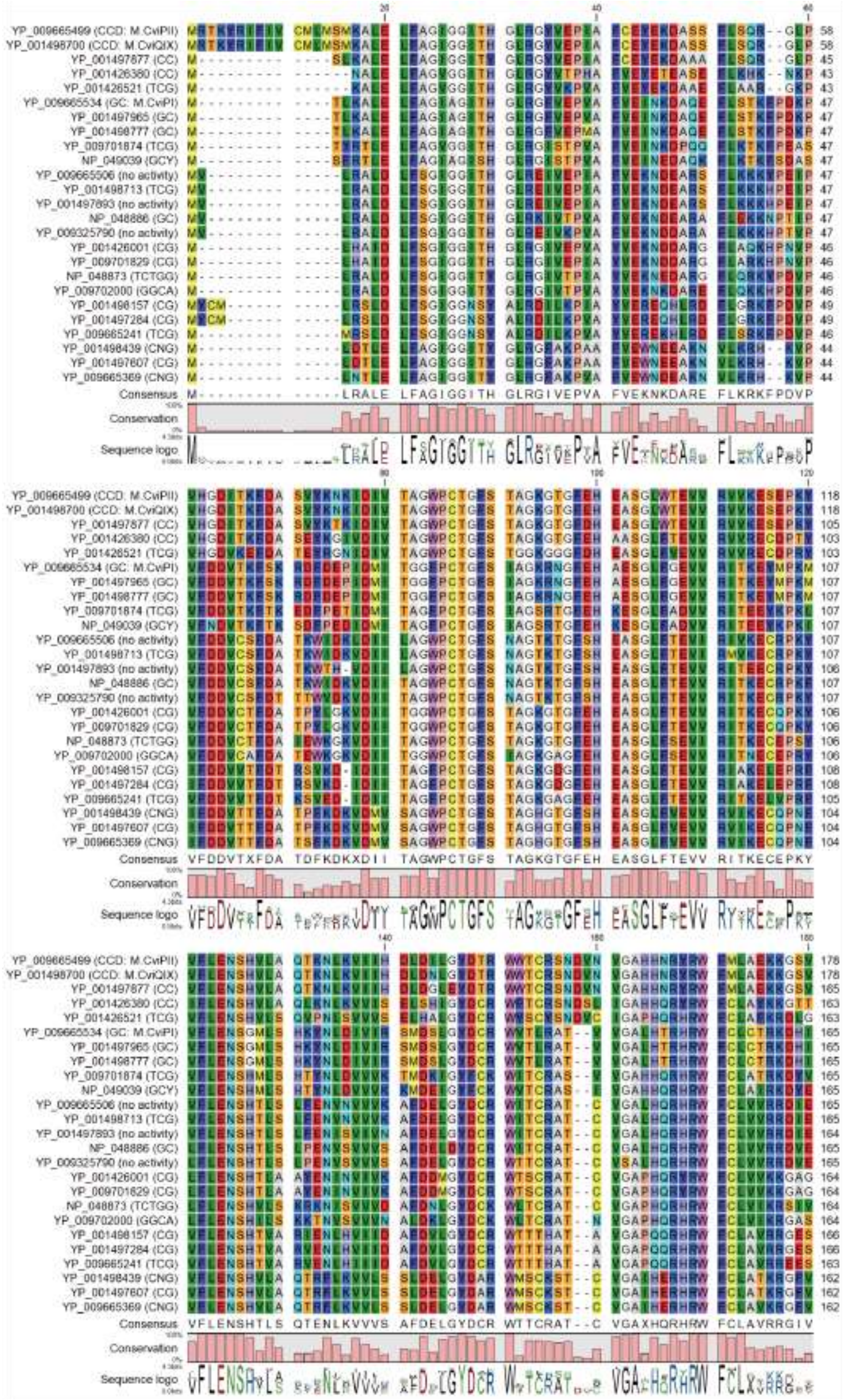

(Continues on the next page)

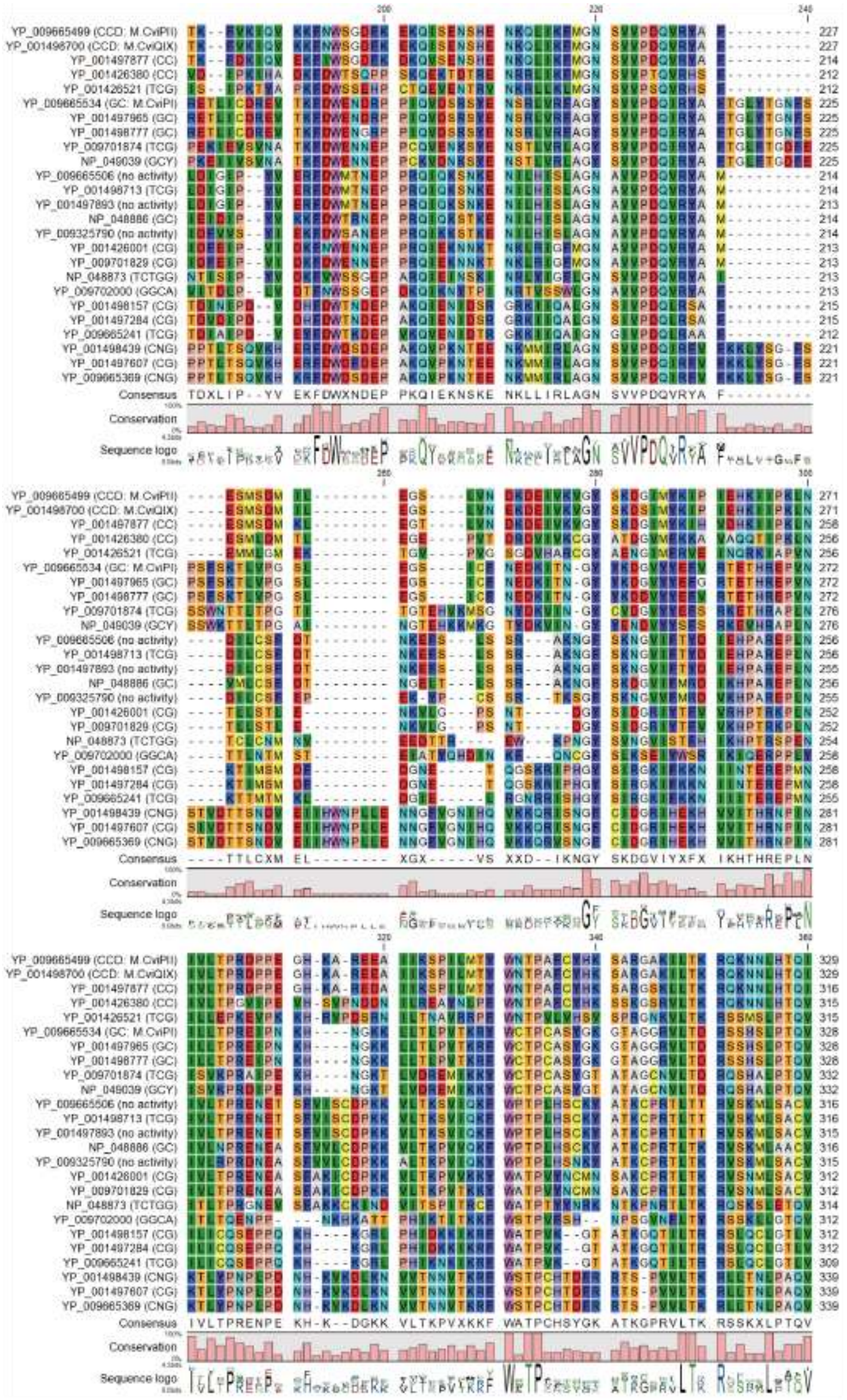


*(Continues on the next page)*

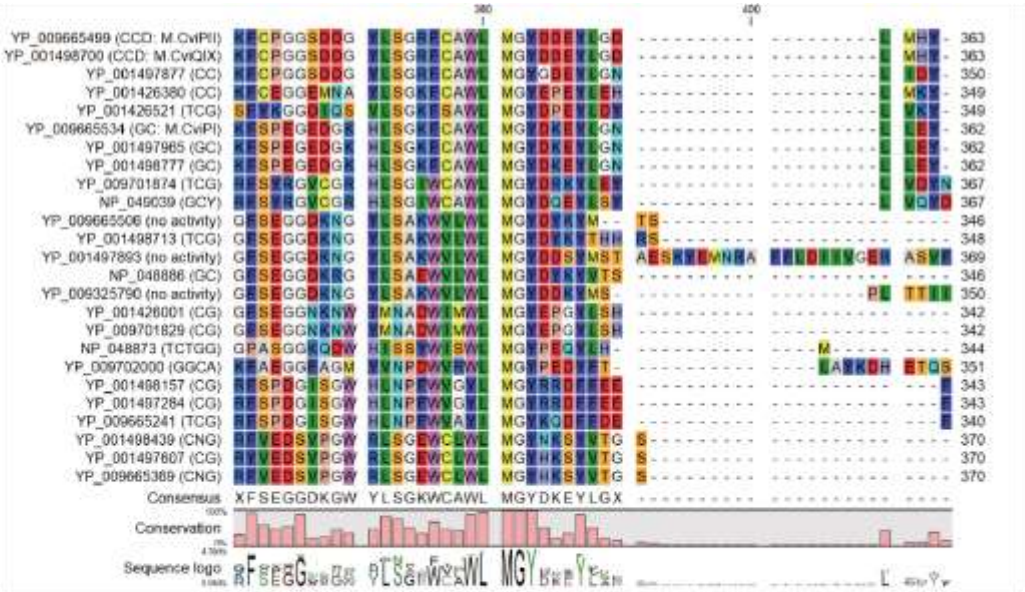

**Supplementary Figure S1 Multiple alignments of the 25 genes.**

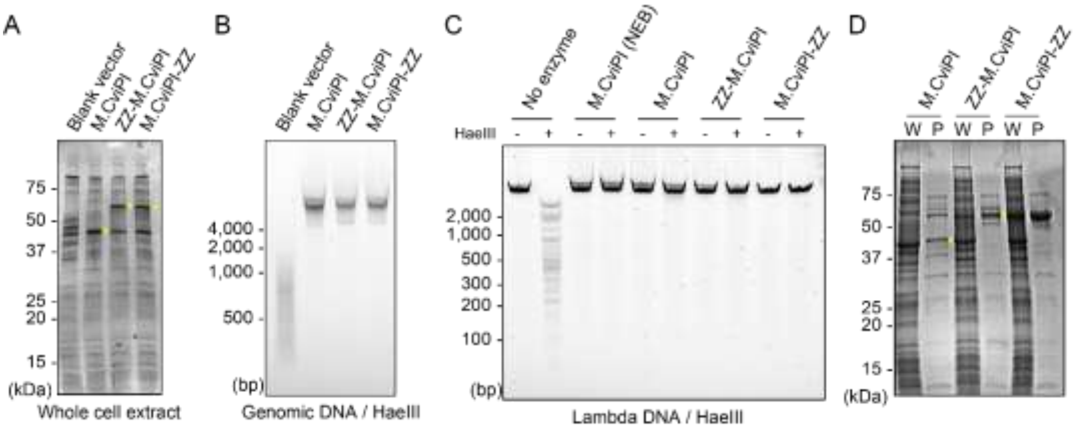


**Supplementary Figure S2 Expression and purification of M.CviPI.**

A. SDS-PAGE of whole-cell extracts of *E. coli* cells with induced expression of indicated constructs. B. Genomic DNA extracted from *E. coli* cells with induced expression of indicated constructs and digested with 5-methylation-sensitive restriction enzyme HaeIII. C. Unmethylated lambda DNA was treated with MTases indicated and digested with HaeIII. D. SDS-PAGE of purified MTases expressed from indicated constructs.

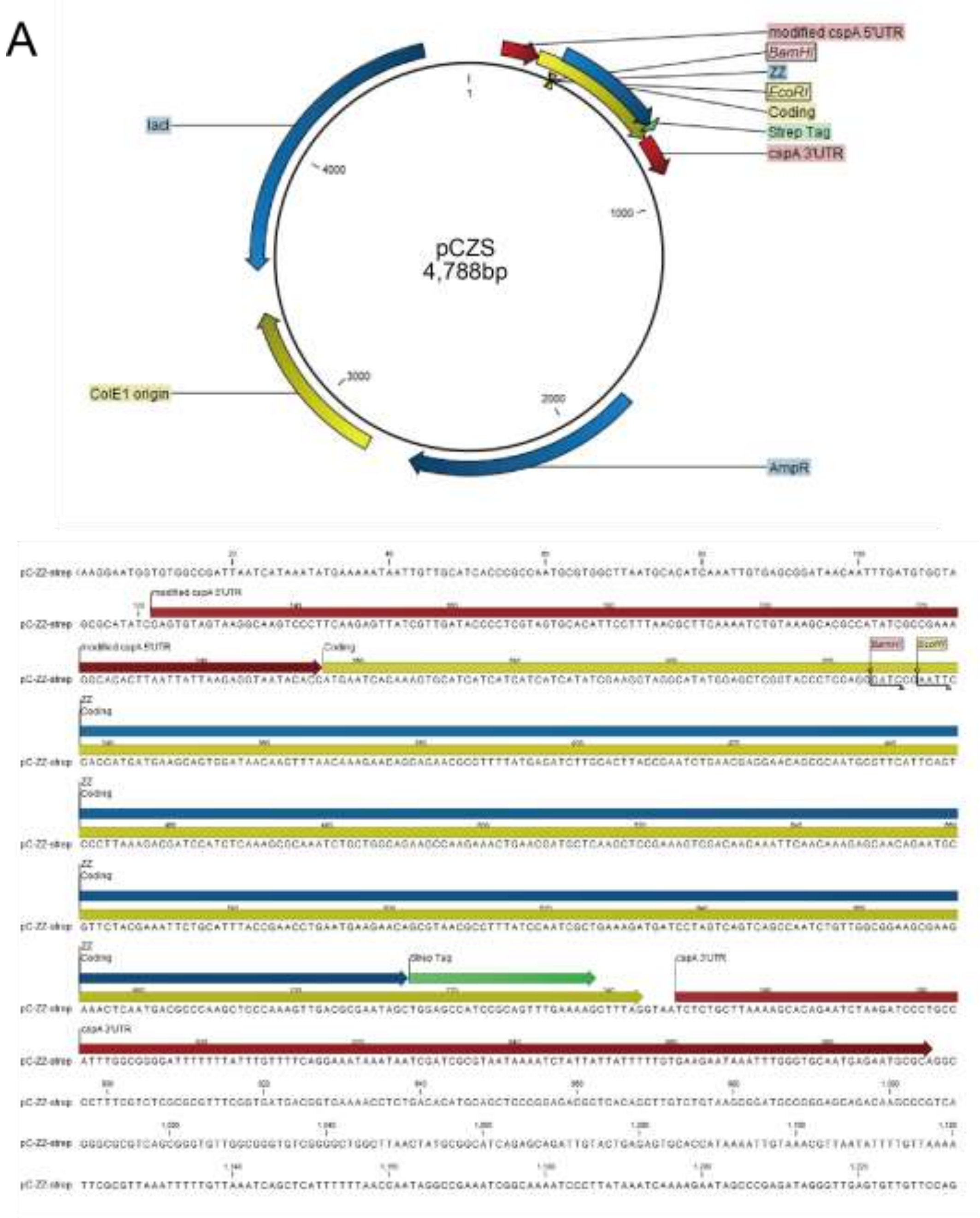


*(Continues on the next page)*

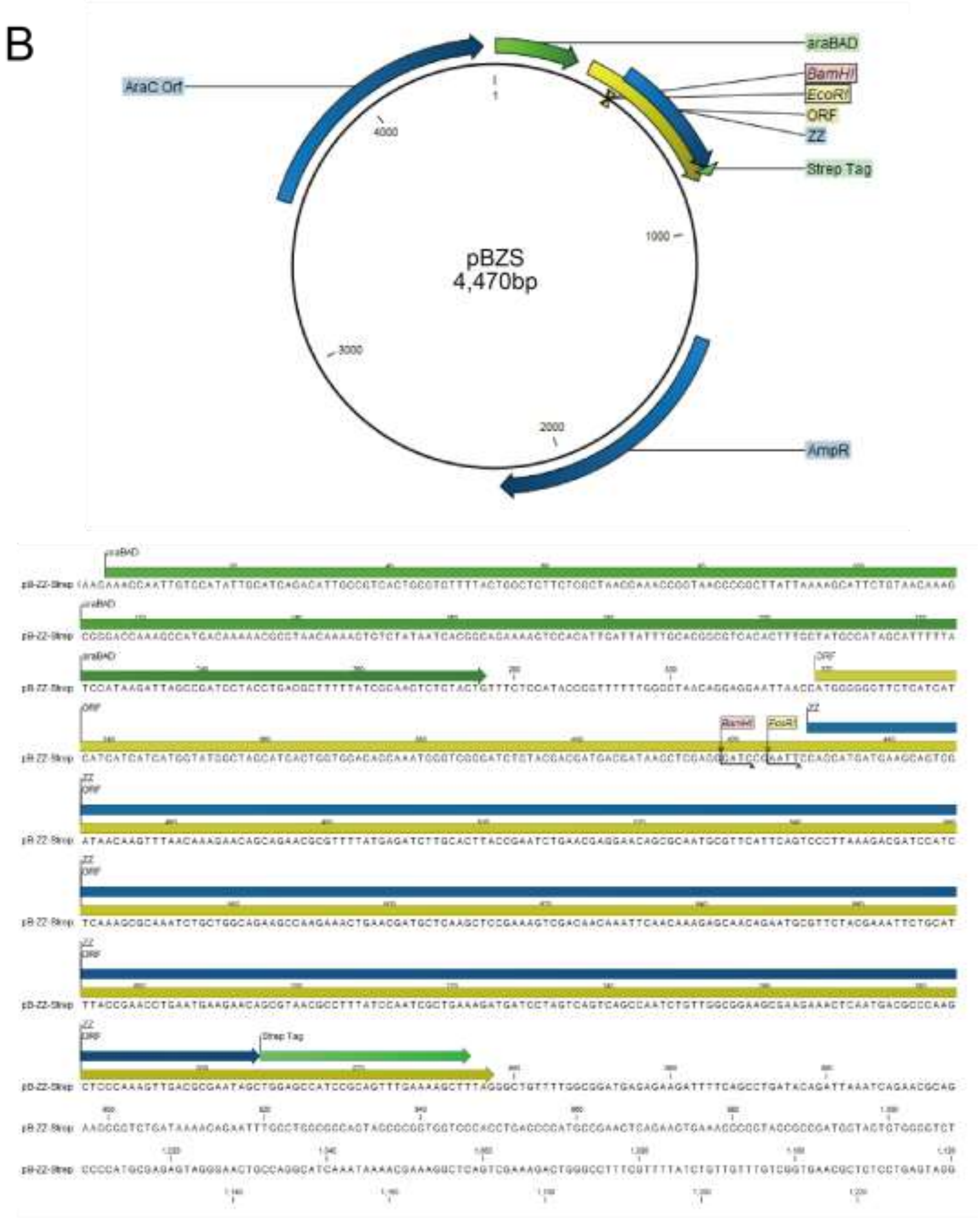


**Supplementary Figure S3 Schemes of expression vectors used.**

The structure of expression vectors (top panels) and sequences of cloning sites (bottom panels) are shown. Candidate MTase-encoding genes were inserted at the BamHI-EcoRI site. Two expression vectors under the control of the cold-shock promoter (A) and arabinose inducible promoter (B) were used.

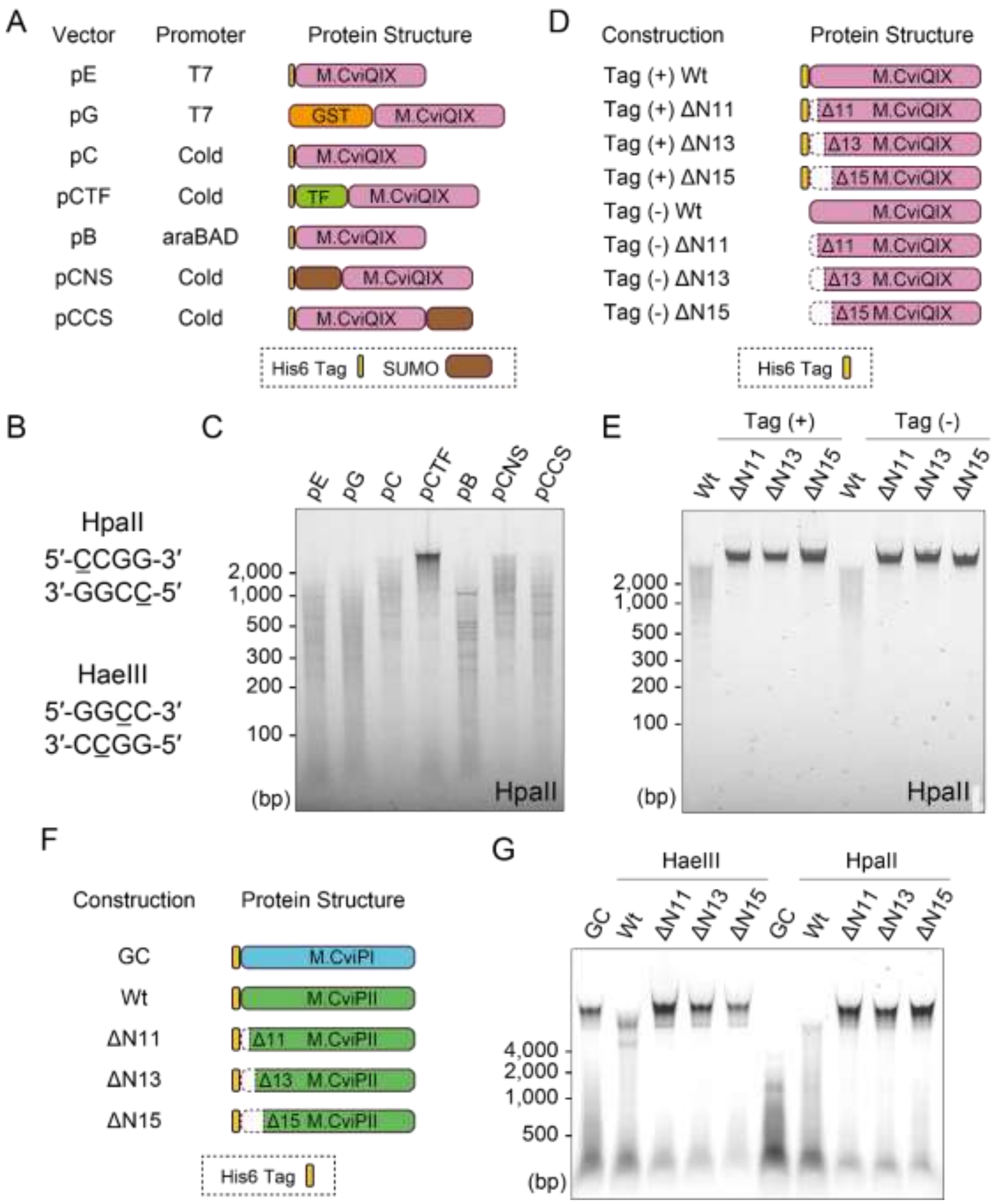


**Supplementary Figure S4 Expression of M.CviQIX and M.CviPII from several constructs.**

A. Schemes of constructions used for the expression of M.CviQIX. B. Recognition sequences of restriction enzymes used to investigate methylation states of genomic DNA. When the underlined C is methylated, the activity of the enzymes is inhibited. C. Analysis of methylation states of the genomic DNA extracted from *E. coli* cells with induced expression of M.CviQIX from the constructions shown in A. D. Schemes of N-terminal deletion mutants of M.CviQIX. A cold shock promoter-based constructions were used. E. Digestion assay of genomic DNA extracted from *E coli* cells harboring the construct shown in D. F. Deletion mutants of M.CviPII were compared. G. Digestion of genomic DNA extracted from *E. coli* cells harboring the constructs in F.

| Construction Name | PWM-M | M-Logo |
| --- | --- | --- |
| pBZS-M.CviPI | 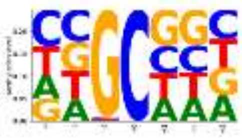 | 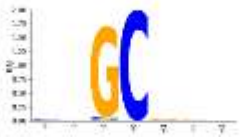 |
| pBZS-M.CviQIX (NDel15) | 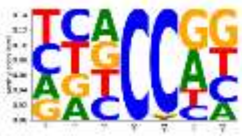 | 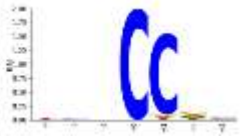 |
| pBZS-NP_048873 | 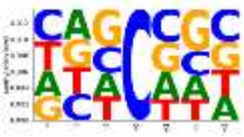 | 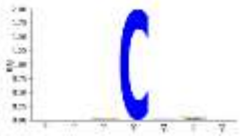 |
| pBZS-NP_048886 | 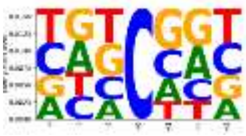 | 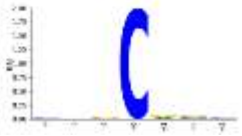 |
| pBZS-NP_049039 | 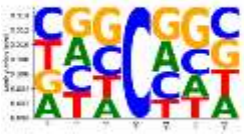 | 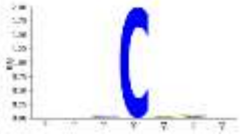 |
| pBZS-YP_001426001 | 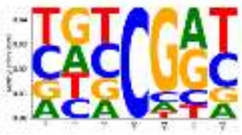 | 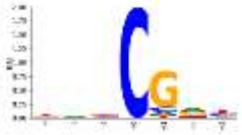 |
| pBZS-YP_001426380 | 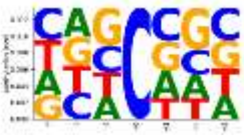 | 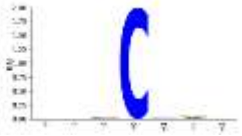 |
| pBZS-YP_001426521 | 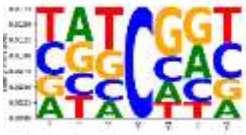 | 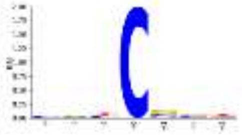 |
| pBZS-YP_001497284 | 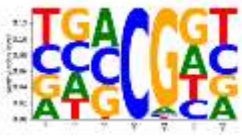 | 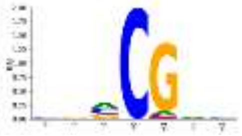 |
| pBZS-YP_001497607 | 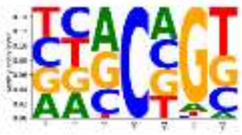 | 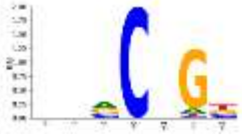 |

*(Continues on the next page)*

| Construction Name | PWM-M | M-Logo |
|---|---|---|
| pBZS-YP_001497877 | 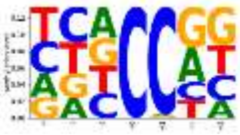 | 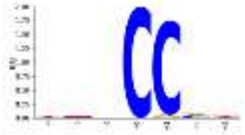 |
| pBZS-YP_001497893 | 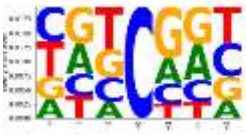 | 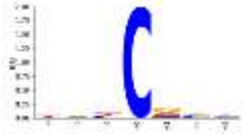 |
| pBZS-YP_001497965 | 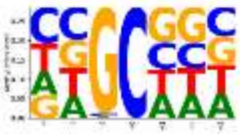 | 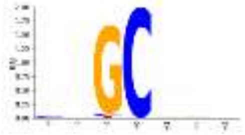 |
| pBZS-YP001498157 | 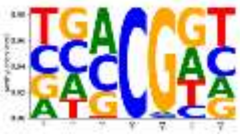 | 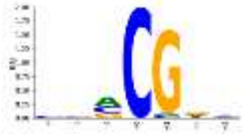 |
| pBZS-YP001498439 | 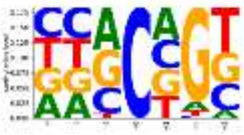 | 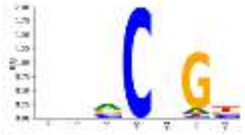 |
| pBZS-YP_001498713 | 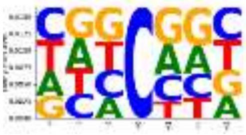 | 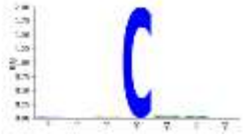 |
| pBZS-YP_001498777 | 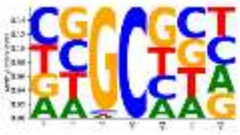 | 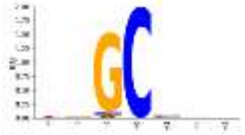 |
| pBZS-YP_009325790 | 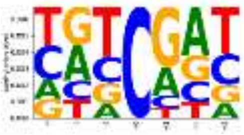 | 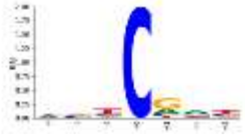 |
| pBZS-YP009665241 | 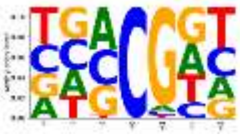 | 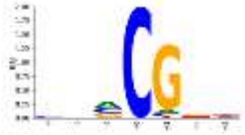 |
| pBZS-YP009665369 | 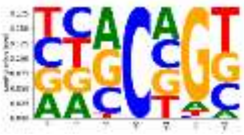 | 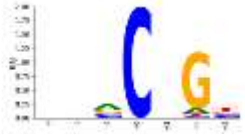 |

*(Continues on the next page)*

| Construction Name | PWM-M | M-Logo |
|---|---|---|
| pBZS-YP_009665506 | 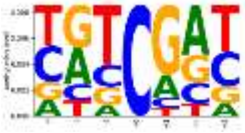 | 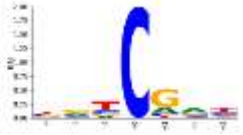 |
| pBZS-YP_009701829 | 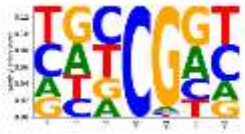 | 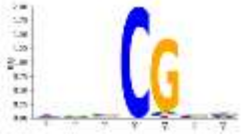 |
| pBZS-YP_009701874 | 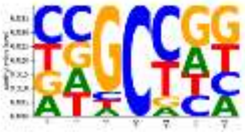 | 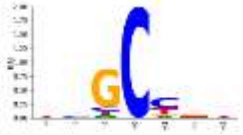 |
| pBZS-YP_009702000 | 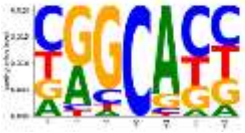 | 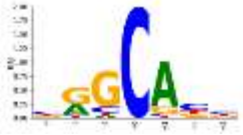 |
| pCZS-M.CviPI | 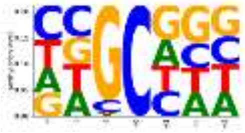 | 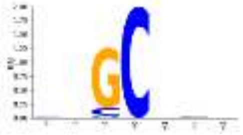 |
| pCZS-M.CviQIX (NDel15) | 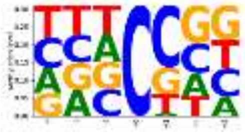 | 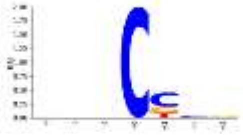 |
| pCZS-NP_048873 | 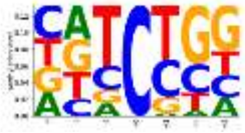 | 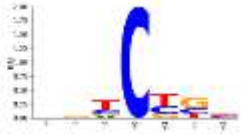 |
| pCZS-NP_048886 | 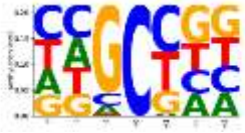 | 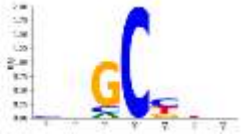 |
| pCZS-NP_049039 | 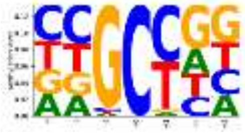 | 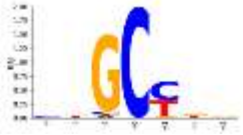 |
| pCZS-YP_001426001 | 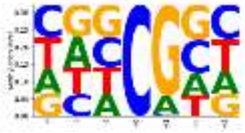 | 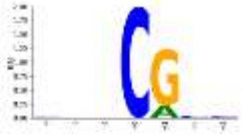 |

*(Continues on the next page)*

| Construction Name | PWM-M | M-Logo |
|---|---|---|
| pCZS-YP_001426380 | 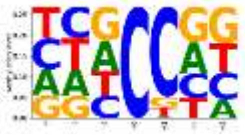 | 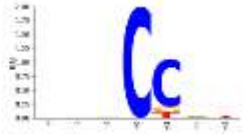 |
| pCZS-YP_001426521 | 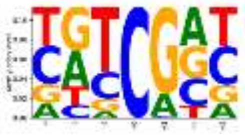 | 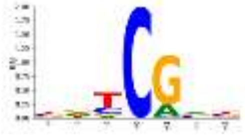 |
| pCZS-YP_001497607 | 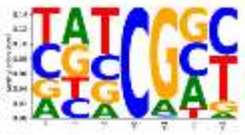 | 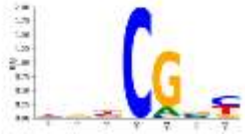 |
| pCZS-YP_001498713 | 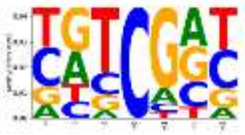 | 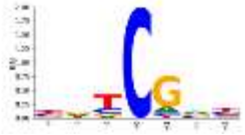 |
| pCZS-YP_009325790 | 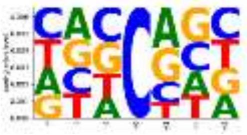 | 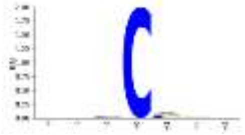 |
| pCZS-YP_009665241 | 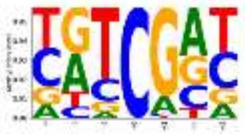 | 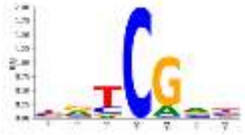 |
| pCZS-YP009665506 | 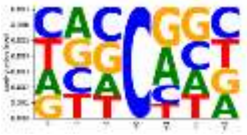 | 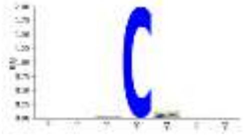 |
| pCZS-YP009701874 | 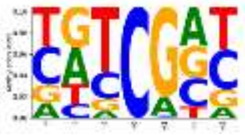 | 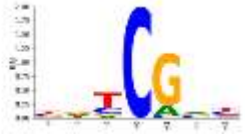 |

**Supplementary Figure S5 PWM-M and M-Logo for MTases.**

PWM-M and M-Logo drawn for methylome data obtained with whole-genome bisulfite sequencing (WGBS) of genomic DNA extracted from *E. coli* cells expressing indicated constructs are shown.

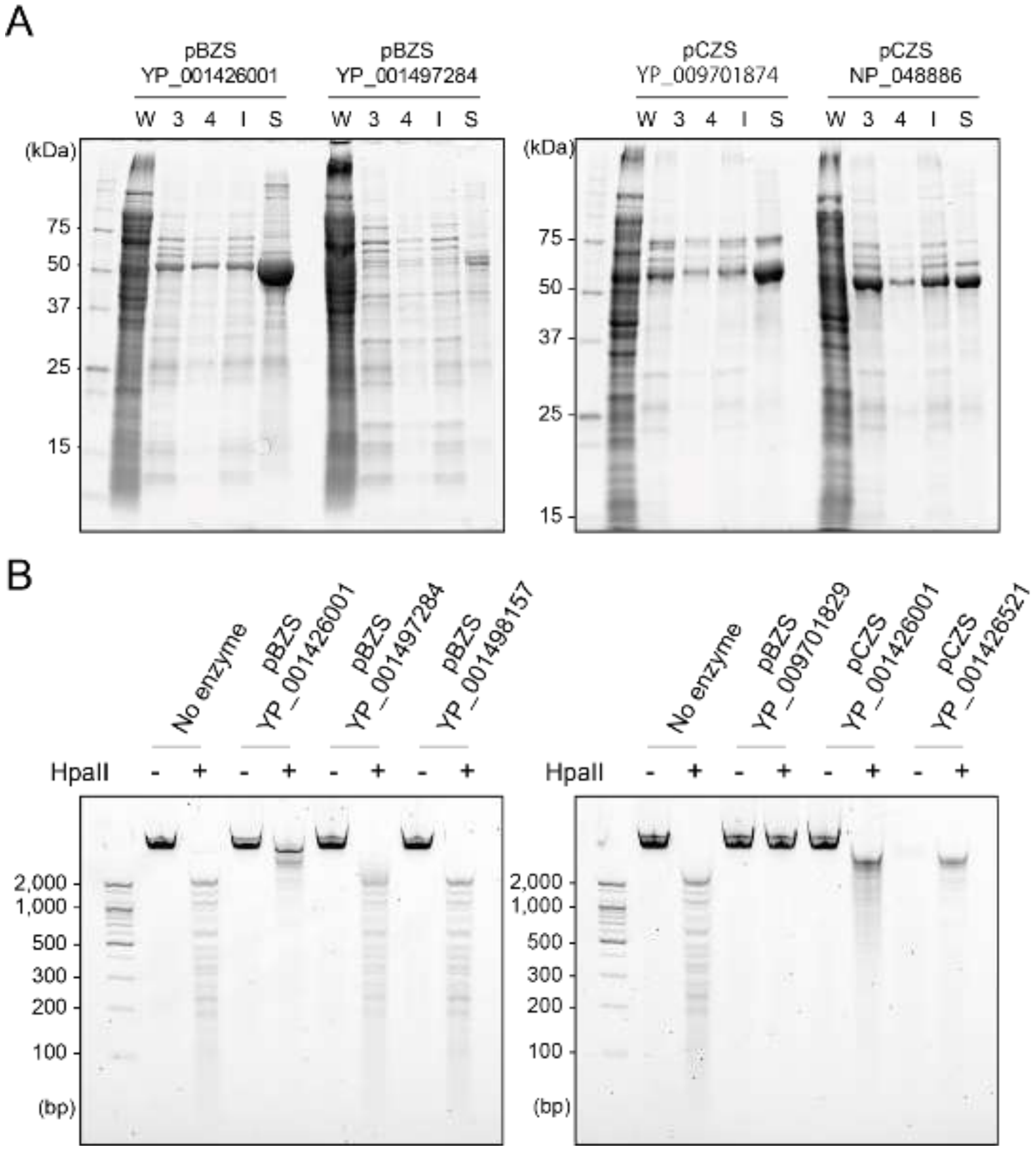


**Supplementary Figure S6 Purification of MTases and their activities.**

A. Representative SDS-PAGE images of protein purifications. W: whole-cell extract; 3 and 4: 3rd and 4th fractions of HisTrap column purification, respectively; I: input of StrepTrap purification; S: purified protein. B. Analysis of MTase activity of purified proteins. Unmethylated lambda DNA was treated with the purified MTases, digested with the methylation-sensitive restriction enzyme HpaII, and analyzed on E-Gel Ex.

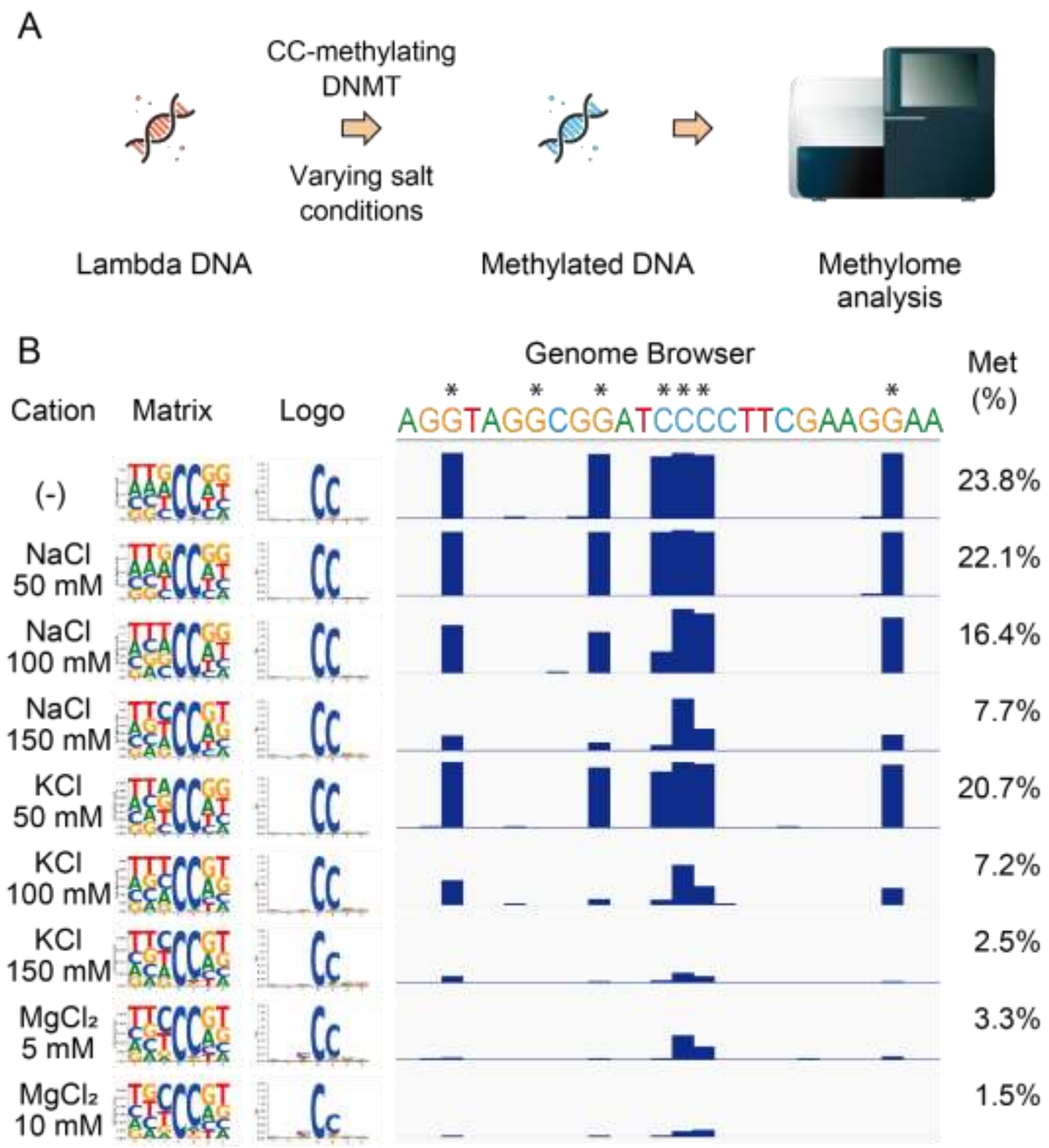


**Supplementary Figure S7 Investigation of optimum salt concentrations for the CCMT.**

A. Strategy used for the analysis. Unmethylated lambda DNA served as substrate for CCMT, and then WGBS was conducted. B. Methylation levels observed at different salt concentrations. The assays were performed in 50-µL reactions that contain 50 mM Tris-HCl (pH8.5), 160 µM S-adenosylmethionine, salts indicated, 1 µg of unmethylated lambda DNA, and 1 µg of purified CCMT. After incubation at 37 °C for 1 h, DNA was purified and used for WGBS. PMM-M (Matrix), M-Logo (Logo), a representative genome browser shot showing methylation levels for individual cytosines, and mean methylation levels of entire lambda DNA are shown for conditions investigated. The asterisks at the top of browser shot indicate the position to be methylated by the CCMT.

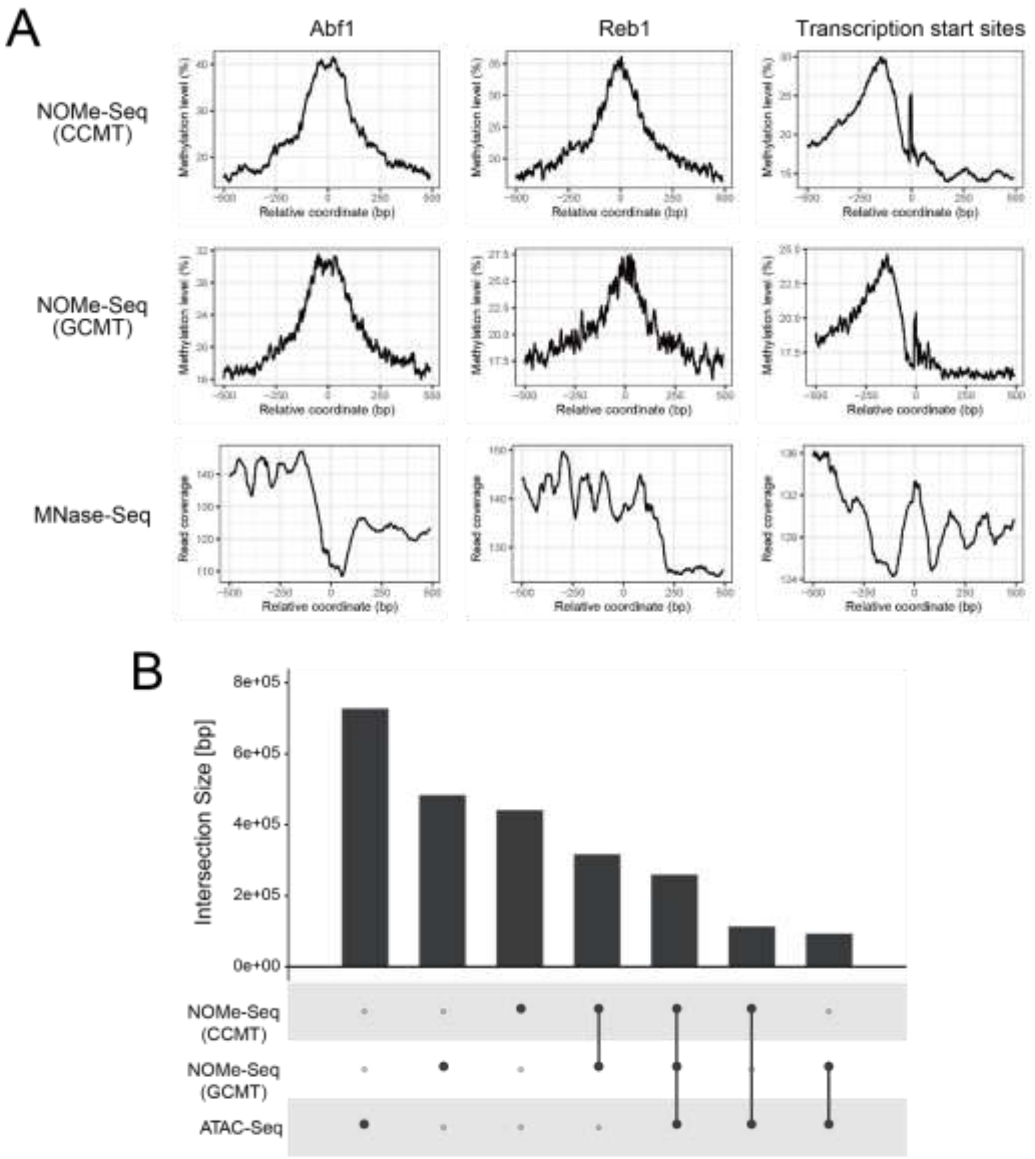


**Supplementary Figure S8 Analysis of fixed budding yeast nuclei with GCMT- and CCMT-based NOMe-Seq.**

A. Aggregation plots for methylation levels of GCMT-, CCMT-based NOMe-Seq, and mapped read coverage of MNase-seq. X axis shows the relative coordinates from the genomic features indicated and the Y axis shows the mean DNA methylation levels (NOMe-Seq) and mapped read coverage (MNase-seq). B. Upset plot showing the overlapping peaks identified with GCMT- and CCMT-based NOMe-Seq and ATAC-Seq.

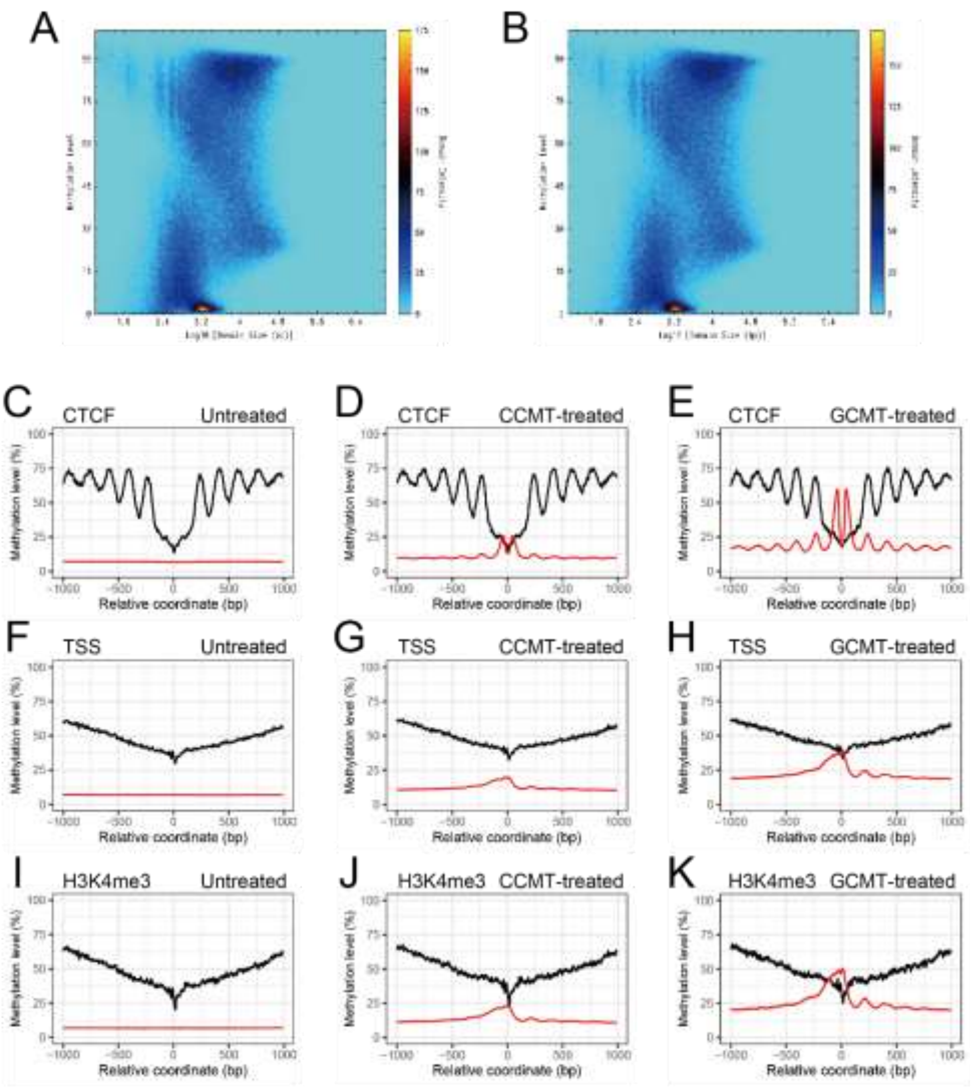


**Supplementary Figure S9 NOMe-Seq analysis of IMR-90 cells.**

A and B. Methylation domain landscape plots for untreated (A) and CCMT-treated (B) IMR-90 cells. The X-axis indicates the domain size, and the Y-axis shows the mean methylation level of the domains. C–K. Aggregation plots centered on the genomic features indicated. The methylation levels of untreated (C, F, and I), CCMT-treated (D, G, and J), and GCMT-treated (E, H, and K) cells are shown. For GCMT-based NOMe-Seq, data from Kelly et al. were used. Black lines indicate the intrinsic methylation levels at CG sites (untreated and CCMT-treated cells) and HCG sites (GCMT-treated cells). Red lines denote the methylation levels induced with MTases at CC sites (untreated and

CCMT-treated cells) and GCH sites (GCMT-treated cells).

**Supplementary tables**

**Supplementary Table S1**

Supplementary Table S1 is provided as an excel file.

**Supplementary Table S2 Basic statistics of the methylome data of fixed yeast nuclei.**

| | | | CCMT-treated | GCMT-treated |
|---|---|---|---|---|
| Accession numbers at NCBI GEO | | | GSM6111857 | GSM5761261 |
| Mapping | Read 1 | Number of reads | 5,882,971 | 3,022,479 |
| | | Unique map | 78.6% | 75.0% |
| | | Unmap | 11.7% | 13.3% |
| | Read 2 | Number of reads | 5,882,967 | 3,022,479 |
| | | Unique map | 76.5% | 74.3% |
| | | Unmap | 15.3% | 14.2% |
| Mapped read coverage | | Top | 26.7 | 13.1 |
| | | Bottom | 26.5 | 13.2 |
| | | Both | 53.2 | 26.3 |
| | | N | 26.6 | 13.1 |
| | | C | 26.6 | 12.4 |
| | | CpG | 27.2 | 12.5 |
| | | CHG | 27.4 | 12.3 |
| | | CHH | 26.7 | 12.6 |
| Methylation level | | C | 2.7% | 2.7% |
| | | CpG | 2,9% | 2.9% |
| | | CHG | 2.5% | 2.5% |
| | | CHH | 2.7% | 2.7% |
| Coverage vs read depth | | >=1 | 95.0% | 94.9% |
| | | >=3 | 94.7% | 94.3% |
| | | >=5 | 94.5% | 92.6% |
| | | >=10 | 92.0% | 71.9% |

**Supplementary Table S3 Basic statistics of the methylome data of IMR-90 cells.**

| | | | Control | CCMT-treated | GCMT-treated |
|---|---|---|---|---|---|
| Accession numbers for NCBI GEO | | | GSM5769478 | GSM5769479 | GSM6111856 |
| Mapping | Read 1 | Number of reads | 462,135,990 | 445,678,391 | 447,579,538 |
| | | Unique map | 73.00% | 72.80% | 69.5% |
| | | Unmap | 24.20% | 24.70% | 27.9% |
| | Read 2 | Number of reads | 462,135,988 | 445,678,391 | 447,579,537 |
| | | Unique map | 65.10% | 64.90% | 63.6% |
| | | Unmap | 32.60% | 32.90% | 34.0% |
| Mapped read coverage | | Top | 12.9 | 12.3 | 11.4 |
| | | Bottom | 12.8 | 12.2 | 11.3 |
| | | Both | 25.7 | 24.5 | 22.7 |
| | | N | 12.9 | 12.3 | 11.3 |
| | | C | 14.2 | 13.4 | 12.4 |
| | | CpG | 14.2 | 13.1 | 12.0 |
| | | CHG | 14.4 | 13.6 | 12.8 |
| | | CHH | 14.1 | 13.4 | 12.5 |
| Methylation level | | C | 3.4 | 3.9 | 5.4 |
| | | CpG | 56.2 | 56.2 | 58.0 |
| | | CHG | 1 | 1.2 | 3.1 |
| | | CHH | 0.8 | 1.5 | 2.8 |
| Coverage vs read depth | | >=1 | 86.2% | 86.1% | 85.7% |
| | | >=3 | 84.0% | 83.8% | 82.2% |
| | | >=5 | 80.9% | 80.6% | 76.9% |
| | | >=10 | 64.8% | 63.2% | 55.5% |

### Supplementary information

### Sequences synthesized in the current study

The restriction sites are indicated with underlines.

>M.CviPI
GGATCCATGACCCTGAAGGCCCTGGAGCTGTTCGCCGGCATCGCCGGCATCACCCACGGC
CTGAGAGGCTTCGTGGAGCCCGTGGCCTTCGTGGAGATCAACAAGGACGCCCAGGAGTTC
CTGAGCACCAAGTTCCCCGACAAGCCCGTGTTCGACGACGTGACCAAGTTCAGCAAGAGA
GACTTCGACGAGCCCATCGACATGATCACCGGCGGCTTCCCCTGCACCGGCTTCAGCATC
GCCGGCAAGAGAAACGGCTTCGAGCACGCCGAGAGCGGCCTGTTCGGCGAGGTGGTGAGA
ATCACCAAGGAGTACATGCCCAAGATGGTGTTCCTGGAGAACAGCGGCATGCTGAGCCAC
AAGTACAACCTGGACATCGTGATCAGAAGCATGGACAGCCTGGGCTACGACTGCAGATGG
GTGACCCTGAGAGCCACCGTGGTGGGCGCCCTGCACACCAGACACAGATGGTTCTGCCTG
TGCACCAGAAAGGACCACATCAGAGAGACCCTGATCTGCGACAGAGAGGTGACCAAGTTC
GACTGGGAGAACGACAGACCCCCCATCCAGGTGGACAGCAGAAGCTACGAGAACAGCAGA
CTGGTGAGATTCGCCGGCTACAGCGTGGTGCCCGACCAGATCAGATACGCCTTCACCGGC
CTGTACACCGGCAACTTCAGCCCCAGCTTCAGCAAGACCCTGGTGCCCGGCAGCCTGGAG
GGCAGCATCTGCTTCAACGAGGACAAGATCACCAACGGCTACTACAAGGACGGCGTGTAC
TACGAGTTCGTGAGAACCGAGACCCACAGAGAGCCCGTGAACATCCTGCTGACCCCCAGA
GAGATCCCCAACAAGCACAACGGCAAGAAGCTGCTGACCCTGCCCGTGACCAAGAGATAC
TGGTGCACCCCCTGCGCCAGCTACGGCAAGGGCACCGCCGGCGGCAGAGTGCTGACCGAC
AGAAGCAGCCACAGCCTGCCCACCCAGGTGAAGTTCAGCCCCGAGGGCGAGGACGGCAAG
CACCTGAGCGGCAAGTTCTGCGCCTGGCTGATGGGCTACGACAAGGAGTACCTGGGCAAC
CTGCTGGAGTACGAATTC

>M.CviPII
GGATCCATGCGCACGAAATACCGCATCTTCATCGTCTGCATGCTGATGTCCATGAAAGCC
TTGGAACTATTTGCGGGGATAGGTGGCATTACCCATGGCTTACGTGGCTATGTAGAACCG
ATTGCATTCTGCGAATATGAGAAAGATGCCAGCTCGTTTCTGTCACAACGTGGCTTACCC
GTTCATGGGGACATTACGAAGTTTGACGCCAGTGTGTACAAAAACAAGATTGACATTGTG
ACAGCAGGATGGCCTTGTACTGGGTTTTCGACCGCGGGTAAAGGCACTGGTTTTGAGCAT
GAAGCTAGCGGTTTATGGACAGAAGTGGTCCGAGTAGTGAAAGAGAGCGAACCGAAATAC
GTCTTTTTGGAAAATAGTCACGTTTTGGCGCAAACCAAAAACCTCAAGGTCATCATCCAT
GATCTGGATATCCTTGGCTATGATACGCGTTGGTGGACGTGCAGATCGAATGACGTTAAT
GTCGGTGCACATCACAATCGGTATCGCTGGTTTATGCTCGCAGAGAAAAAGGCTCTGTG
ACCAAATTCGTGAAAATTCAGGTGAAAAAGTTCAATTGGTCTGGTGACTTCAAAGAGAAA
CAGATAAGCGAGAACAGTCACGAAAACAAGCAGCTCATTAAGTTCATGGGAAACTCCGTA
GTTCCGGATCAAGTTCGCTATGCGTTTGAATCGATGAGCGACATGATTCTGGAAGGCAGT
CTGGTGAATGATAAGGATGAGATCGTGAAGGTAGGCTATTCCAAAGATGGCATCATGTAC
AAAATTCCGATTGAACACAAAATCATTCCGAAACTTAACATCGTTCTGACCCCACGTGAT
CCGCCAGAAGGCCATAAAGCCCGTGAAGAAGCGATCATCAAAAGCCCCATTCTGATGACG
TACTGGAATACTCCGGCCTTTTGCTATCACAAATCAGCTCGCGGCGCTAAAATTCTGACC
AAACGTCAGAAAAACAACCTGCATACCCAGATCAAATTCTGTCCTGGTGGTTCTGACGAT
GGTTATCTGTCAGGTCGCTTTTGTGCGTGGCTGATGGGATACGATGATGAATACCTGGGG
GATCTTATGCACTATGAATTC

>M.CviQIX
GGATCCATGCGCACGAAATACCGCATCTTCATCGTCTGCATGCTGATGTCCATGAAAGCC
TTGGAACTATTTGCGGGGATAGGTGGCATTACCCATGGCTTACGTGGCTATGTAGAACCG
ATTGCATTCTGCGAATATGAGAAAGATGCCAGCTCGTTTCTGTCACAACGTGGCTTACCC
GTTCATGGGGACATTACGAAGTTTGACGCCAGTGTGTACAAAAACAAGATTGACATTGTG
ACAGCAGGATGGCCTTGTACTGGGTTTTCGACCGCGGGTAAAGGCACTGGTTTTGAGCAT
GAAGCTAGCGGTTTATGGACAGAAGTGGTCCGAGTAGTGAAAGAGAGCGAACCGAAATAC
GTCTTTTTGGAAAATAGTCACGTTTTGGCGCAAACCAAAAACCTCAAGGTCATCATCCAT
GATCTGGATAACCTTGGCTATGATACGCGTTGGTGGACGTGCAGATCGAATGACGTTAAT
GTCGGTGCACATCACAATCGGTATCGCTGGTTTATGCTCGCAGAGAAAAAAGGCTCTGTG
ACCAAATTCGTGAAAATTCAGGTGAAAAAGTTCAATTGGTCTGGTGACTTCAAAGAGAAA
CAGATAAGCGAGAACAGTCACGAAAACAAGCAGCTCATTAAGTTCATGGGAAACTCCGTA
GTTCCGGATCAAGTTCGCTATGCGTTTGAATCGATGAGCGACATGATTCTGGAAGGCAGT
CTGGTGAATGAAAAGGATGAGATCGTGAAGGTAGGCTATTCCAAAGATAGCATCATGTAC
AAAATTCCGATTGAACACAAAATCATTCCGAAACTTAACATCGTTCTGACCCCACGTGAT
CCGCCAGAAGGCCATAAAGCCCGTGAAGAAGCGATCATCAAAAGCCCCATTCTGATGACG
TACTGGAATACTCCGGCCTTTTGCTATCACAAATCAGCTCGCGGCGCTAAAATTCTGACC
AAACGTCAGAAAAACAACCTGCATACCCAGATCAAATTCTGTCCTGGTGGTTCTGACGAT
GGTTATCTGTCAGGTCGCTTTTGTGCGTGGCTGATGGGATACGATGATGAATACCTGGGG
GATCTTATGCACTATGAATTC

>NP_048873
GGATCCATGCTGCGTGCGCTGGACCTGTTCAGCGGCATCGGTGGCATTACCTACGGCCTG
CGTGGTATCGTTACCCCGATTGCGTATGTGGAGAAGAACGAAGACGCGCGTGGCTTTCTG
CAGCGTAAATACCCGGATGTTCCGGTGTTCGACGATGTTTGCACCTTTGACGCGATCGAG
TGGAAGGGTAAAGTGGATATCATTACCGCGGGTTGGCCGTGCACCGGTTTCAGCACCGCG
GGCAAGGGCACCGGTTTCGAGCACGAAGCGAGCGGTCTGTTTAGCGAGGTTATCCGTATT
ACCAAAGAGTGCGAACCGAGCTACCTGTTCCTGGAAAACAGCCACGTGCTGAGCAAGCGT
AAAAACATCAGCGTGGTTGTGGACGCGTTTGATAACCTGGGCTATGACTGCAAGTGGCTG
ACCTGCCGTGCGACCTGCGTTGGTGCGCCGCACCAACGTCACCGTTGGTTCTGCCTGGTT
ATCAAGCGTAGCATTGTGAACACCATCAGCATTCCGTACGTTGATAAATTTGTTTGGAGC
AGCGGTGAACCGGCGCGTCAGATCGAAATTAACAGCAAAATTAACCGTCTGTACATCGGC
TTCCTGGGTAACAGCGTTGTGCCGGACCAAGTTCGTTATGCGATTACCTGCCTGTGCAAC
ATGAACGTGGAGGAAGATACCACCCGTGAATGGAAGCCGAACGGCTATAGCGTTAACGGT
GTGATTAGCACCTTTCACATCAAACACCCGACCCGTAGCCCGGAGAACATCACCCTGACC
CCGCGTGGTAACGAAGTTAGCTTCGCGAAGAAATGCAAGATCAACGACGTGATTACCAGC
CCGATCACCCGTTGCTTTTGGGCGACCCCGACCTACTATAACCGTAAGAACACCAAACCG
AACCGTACCCTGACCAAGCGTCAGAGCAAAAGCCTGGAGACCCAGGTTGGTCCGGCGAGC
GGTGGCAAACAGGATTGGCACATTAGCAGCTATTGGATCAGCTGGCTGATGGGTTACCCG
GAACAATATCTGCACATGGAATTC

>NP_048886
CTCGAGGGATCCATGGTACTTCGTGCGTTAGATCTGTTCAGTGGAATTGGCGGTATTACC
CATGGACTCCGCAAAATCGTGACACCAGTGGCATTTGTGGAAAAGAACGATGAAGCTCGT
GCCTTTCTGGAGAAGAAAAACCCGACTATTCCTGTGTTTGACGATGTATGCTCATTTGAT
GCGACCAAATGGATTGACAAAGTCGACATTATTACGGCAGGTTGGCCTTGTACGGGCTTT
TCCAATGCTGGTACCAAAACGGGGTTTTCGCATGAAGCGAGCGGTCTGTTTACCGAGGTA
GTTCGCATTACCAAAGAATGTCGGCCGAAATTCCTGTTCCTGGAGAATAGCCACACTCTG
TCCTTACCGGAAAACGTGTCAGTTGTCGTTTCTGCCTTTGATGAGCTGGATTATGACTGT
CGCTGGATTACCTGCCGTGCAACGTGCGTTGGTGCCCTCCATCAACGCCATCGCTGGTTC
TGCTTAGTTGTGCGTCGAGATGTGGAAATCGAAATTGACATACCGTACGTGAAAAAGTTT
GATTGGACACGCAATGAACCGCCCAGGCAAATCCAGAAAAGCACCAAAGAGAACATCCTG
CATATCAGCTTAGCGGGCAATGCCGTTGTCCCGGATCAGGTACGCTATGCCATGGTCATG
CTGTGCTCTTTTGACACCAATGGCGAACTTACGCTCAGTAGCCGTGCGAAAAACGGCTTT
TCGAAAGATGGCGTCATCTTCATGCGTGATATCAAACACCCAGCACGCGAACCGCTGAAC
ATCGTCCTGAATCCCAGAGAGAACGAAGCCTCATTCGTGGTTCTGTGCGATCCGAAAAAA
GTGCTGACCAAACCGGTTGTCCAGAAGTTCTGGCCTACACCATTGCACAGCTGCAAATAC
GCGACTAAGTGTCCGCGTACTTTGACCAAACGGGTGTCGAAAATGCTTGCTGCGTGTGTT
GGCTTCTCCGAAGGTGGGGATAAACGTGGCTATCTGTCTGCGGAATGGGTATTGTGGCTA
ATGGGGTATGACTACAAGTATGTGACGAGTGAATTCAAGCTT

>NP_049039
GGATCCATGTCATTTAGGACATTAGAACTATTCGCTGGCATCGCCGGTATCAGCCACGGC
CTGCGTGGTATTTCCACCCCGGTTGCCTTCGTGGAAATCAACGAAGATGCGCAAAAATTC
CTGAAAACGAAGTTCTCAGACGCGAGCGTGTTCAACGACGTGACCAAATTTACGAAGAGC
GACTTCCCGGAGGACATCGATATGATTACTGCGGGTTTTCCGTGTACCGGCTTCTCGATC
GCTGGCAGCCGTACCGGCTTCGAGCACAAAGAGTCCGGTCTGTTTGCTGATGTGGTGCGT
ATTACCGAGGAATATAAACCGAAGATCGTGTTTTTGGAGAACTCGCACATGCTGAGCCAC
ACCTACAATCTGGATGTAGTCGTCAAAAAGATGGATGAAATTGGTTACTTCTGCAAGTGG
GTTACCTGTCGTGCGAGCATTGTTGGTGCACATCATCAGCGTCACCGCTGGTTTTGCCTG
GCGATCCGCAAGGACTATGAACCGAAAGAGATCATCGTTAGCGTGAATGCCACCAAGTTC
GATTGGGAAAACAACGAGCCGCCTTGCAAGGTTGATAATAAATCTTACGAGAACAGCACA
TTGGTACGCCTGGCTGGCTATAGCGTGGTTCCGGATCAAATTCGTTATGCGTTTACCGGC
CTCTTCACCGGTGATTTTGAATCCAGCTGGAAAACGACTTTAACCCCAGGTGCGATTAAC
GGTACTGAGCACAAGAAGATGAAAGGTACGTACGACAAGGTGATTAACGGCTACTACGAG
AACGACGTGTATTACTCCTTTTCCCGCAAAGAGGTTCATAGAGCACCGCTGAATATTTCT
GTCAAGCCGCGTGACATTCCGGAAAAACACAACGGTAAAACTTTGGTCGACCGTGAAATG
ATCAAAAAGTATTGGTGCACCCCGTGCGCGAGCTACGGCACGGCTACCGCGGGTTGCAAT
GTTCTGACCGATCGTCAGAGCCATGCACTGCCGACCCAGGTTCGTTTTTCTTACCGCGGA
GTTTGTGGTCGTCATCTGAGCGGCATCTGGTGCGCATGGCTGATGGGTTACGACCAAGAA
TACCTAAGCTATTTGGTGCAGTATGACGAATTC

>YP_001426001
GGATCCATGCTGCACGCGATCGACCTGTTCAGCGGCATCGGTGGCATTACCCACGGCCTG
CGTGGTATTGTGGAGCCGGTTGCGTACGTGGAGAAGAACGACGATGCGCGTGGTTTTCTG
GCGCAGAAGCACCCGAACGTGCCGGTTTTCGACGATGTTTGCACCTTTGACGCGACCCCG
TATCTGGGCAAAGTGGATATCATTACCGGTGGCTGGCCGTGCACCGGTTTCAGCACCGCG
GGCAAGGGCACCGGTTTCGAGCACGAAGCGAGCGGTCTGTTTACCGAGGTGGTTCGTATC
ACCAAAGAATGCCAGCCGAAGTACCTGTTCCTGGAGAACAGCCACACCCTGGCGGCGTAT
GAAAACATCAACGTGATTGTTAAAGCGTTTGACGATATGGGCTACGACTGCCGTTGGACC
AGCTGCCGTGCGACCTGCGTTGGTGCGCCGCACCAACGTTATCGTTGGTTCTGCCTGGTG
GTTAAGAAAGGCGCGGGTATTGACTTCGAGATCCCGGTGATTGATAAGTTTAACTGGGAG
AACAACGAACCGCCGCGTCAGATCGAAAAGAACAACAAAACCAACAAGCTGCGTATTGGC
TTTATGGGTAACGCGGTGGTTCCGGATCAAGTTCGTTACGCGATGACCCTGCTGAGCACC
CTGGAGAACAAAGTGCTGGGCCCGAGCAACACCGACGGCTACAGCATCGATGGTCGTATT
TATACCTTCGTGGTTCGTCACCCGACCCGTAAGCCGCTGAACATCGTTCTGACCCCGCGT
GAGAACGAAGCGAGCTTTGCGAAAATTTGCGACCCGAAGAAAGTTCTGACCAAGCCGGTG
GTTAAGAAATACTGGGCGACCCCGGTGTATAACTGCATGAACAGCGCGAAATGCCCGCGT
ACCCTGACCAAGCGTGTTAGCAACATGCTGAGCGCGTGCGTGGGTTTTAGCGAGGGTGGC
AACAAAAACTGGTATATGAACGCGGATTGGATCATGTGGCTGATGGGCTACGAACCGGGT
TATCTGAGCCACGAATTC

>YP_001426380
GGATCCATGAACGCGCTGGAACTGTTTGCGGGTGTGGGTGGCATCACCCATGGTCTGCGT
GGTTACGTGACCCCGCACGCGTTCGTTGAGTATGAAACCGAGGCGAGCGAATTTCTGAAG
CACAAAAACAAGCCGGTTCACGGCGACATTACCAAATTCGATGCGAGCGAGTACAAGGGT
ATCGTGGACATTGTTACCGCGGGCTGGCCGTGCACCGGTTTTAGCACCGCGGGCAAGGGT
ACCGGTTTTGAGCATGCGGCGAGCGGTCTGTTTACCGAAGTGGTTCGTGTGGTTCGTGAG
TGCGATCCGACCTACATCTTCCTGGAAAACAGCCACGTGCTGGCGCAGCTGAAAAACCTG
AAGGTGGTTATCAGCGAGCTGAGCCACATTGGCTATGACTGCCGTTGGTTCACCTGCCGT
AGCAACGATAGCCTGATTGGTGCGCACCACCAACGTTACCGTTGGTTTTGCCTGGCGTAT
AAGAAAGGCACCACCGTTGACATCCCGAAAATTCACGCGGACAAGTTTGATTGGACCAGC
CAGCCGCCGAGCAAACAAGAAAAGACCGATACCCGTGAGAACCGTCGTCTGATCAAGTTC
ATGGGTAACAGCGTGGTTCCGACCCAAGTGCGTCACAGCTTTGAAAGCATGCTGGACATG
ACCCTGGAAGGCGAGCCGGTTACCGACCGTGATGTGATTGTTAAATGCGGCTATGCGACC
GATGGTGTGATGTTCAAGAAAGCGGTTGCGCAGCAAACCATCCCGAAGCTGAACATTGTG
CTGACCCCGGGTGTTATCCCGGAAGTGCACAGCGTTCCGAACGACGATAACATTCTGCGT
GAGGCGTACAACCTGCCGTTCTGGAACACCCCGGCGTTTTGCTATCACAAAAGCAGCAAG
GGCAGCCGTGTGCTGACCAAACGTCAGAAGAACAACCTGCACACCCAAGTTAAATTCTGC
GAAGGTGGCGAGATGAACGCGTACCTGAGCGGCAAGTTTTGCGCGTGGCTGATGGGTTAC
GAACCGGAGTATCTGGAGCACCTGATGAAATATGAATTC

>YP_001426521
GGATCCATGAAGGCGCTGGAACTGTTCGCGGGTGTGGGTGGCATTACCCACGGCCTGCGT
GGTTACGTTAAGCCGGTGGCGTTCGTTGAGTATGAAAAAGATGCGGCGGAGTTTCTGGCG
GCGCGTGGCAAGCCGGTGCACGGTGACGTTAAAGAGTTCGATGCGACCGAATACCGTGGC
AACATCGATATTGTTACCGCGGGCTGGCCGTGCACCGGTTTTAGCACCGGTGGCAAAGGT
GGCGGTTTCGACCACGAGGCGAGCGGTCTGTTTGTGGAAGTGGTTCGTGTGGTTCGTGAG
TGCGATCCGCGTTACGTTTTCCTGGAAAACAGCCACGTGCTGAGCCAGGTTCCGAACCTG
AGCGTGGTTGTGAGCGAGCTGCACGCGCTGGGCTATGACTGCCGTTGGTACAGCTGCTAT
AGCAACGATGTGTGCATCGGTGCGCCGCACCAACGTCACCGTTGGTTCTGCCTGGCGTTT
AAGCGTGACCTGGGCATCAGCATTCCGAAGACCTACGCGCCGAAATTTGATTGGAGCAGC
GAACACCCGTGCACCCAGGAAGTGGAAAACACCCGTGTTAACAAGCGTCTGCTGAAACTG
ATGGGTAACAGCGTTGTGCCGAGCCAAGTGCGTCACAGCTTCGAGATGATGCTGGGCATG
GAAAAAACCGGTGTTCCGGTTGGCAGCGGTGATGTTCATGCGCGTTGCGGCTATGCGGAA
AACGGTATTATGTTTCGTGTGGAGATCAACCAGCGTAAGATTGCGCCGGTTAACATCCTG
CTGGAGCCGAAAGAAGTGCCGAAGAAACACCGTGTTCCGGATAGCCGTAACATCCTGACC
AACGCGGTGCGTCGTCCGTTTTGGAACACCCCGGTTCTGGTGCACAGCGTGAGCCCGCGT
GGTAACAAGGTTCTGACCAAACGTAGCAGCATGAGCCTGCCGACCCAAGTGAGCTTCTAT
AAGGGCGGTGACATTCAAAGCGTTCTGAGCGGCAAATTTAGCGCGTGGCTGATGGGTTAC
GACCCGGAGTACCTGGATTATCTGGTTAAGTATGAATTC

>YP_001497284
GGATCCATGTATTGTATGTTGAGGTCATTAGATCTATTCAGCGGCATCGGTGGTAACTCC
TATGCTTTACGCGACATCCTGAAACCGGTTGCGTATGTGGAGCGCGAGCAGCATCTGCGC
GATTTCCTGGGTCGGAAGTTTCCGGACGTCCCGATTTTCGACGACGTTGTCACCTTTGAT
ACCAGATCCGTGAAGGACATCGATATTATCACCGCGGGCTTCCCGTGCACCGGTTTTTCC
ACGGCTGGCAAGGGCGATGGATTCGAGCACGAGGCGAGCGGTCTGTTTACCGAGGTGGTG
CGCATCGCCAAAGAACTGGAACCGCGTTTTGTTTTTTTGGAGAACTCACATACCGTTGCC
CGTGTCGAGAACCTGCACGTGATTATCGACGCCTTCGATGTGCTGGGCTACGACTGTCGT
TGGACGACCACCCATGCAACTGCGGTGGGTGCGCCTCAACAACGCCATCGTTGGTTCTGC
TTGGCTGTGCGTCGTGGTGAAAGCACCGATGTGGACATCCCGGACGTTGACCACTTTGAT
TGGACCAATGATGAACCAGCGAAGCAGATTGAAAACATTGACAGCCGTGGTCGTAAGATC
ATCCAGGCGCTCGGCAACTCTATCGTTCCGGACCAGCTGAGAAGCGCATTCAAGACCATT
ATGAGCATGGATTTTGACGGCAATGAAACCCAGGGTAGCAAACGCATCCCGCACGGCTAC
TCGATCCGCGGTAAAATCTTCAAGAAGAACATCATTAACACCGAGCGTGAACCGATGAAT
ATCCTGATTTGCCAATCTGAACCACCGCAAAAACACAAAGGCCGTCTGCCGCACATTGAT
AAAAAGATTAAACGTTTCTGGGCAACTCCGGTTAAGGGCACGGCGACGAAAGGTCAGACC
ATTTTGACCCGTCGCAGCCTGCAATGTCTGGGTACATTGGTTCGTTTCTCTCCGGATGGT
ATTAGCGGCTGGCATTTGAATCCGTTTTGGGTTGGCTACCTGATGGGTTATCGTCGTGAC
TTCTTCGAGGAATTTGAATTC

>YP_001497607
GGATCCATGCTGGATACCCTGGAGCTGTTCGCGGGTATCGGTGGCATTACCTATGGTCTG
CGTGGTTTCGCGAAGCCGGCGGCGTTTGTTGAATGGAACGAGGAAGCGAAAAACGTGCTG
AAGCGTCACAAAGTTCCGATCTTCGACGATGTGACCACCTTCGACGCGACCCCGTTTAAG
GATAAAGTGGATATGGTTAGCGCGGGTTGGCCGTGCACCGGTTTTAGCACCGCGGGTCAT
GGTACCGGTTTCAGCCATGAGGCGAGCGGCCTGTTTGTGGAAGTGGTTCGTGTTATCAAG
GAGTGCCAGCCGAACTTCGTTTTTCTGGAAAACAGCCACGTGCTGGCGCAAACCCGTTTC
CTGAAGGTGGTTCTGAGCAGCCTGGATGAGCTGGGCTACGACGCGCGTTGGATGAGCTGC
AAAAGCACCTGCGTGGGTGCGATTCACGAACGTCACCGTTGGTTCTGCCTGGCGACCAAG
CGTGGTTTTGTTCCGCCGACCCTGACCAGCCAGGTGAAACACGAACGTTTCGACTGGGAT
TTTGACGAGCCGGCGAAGCAAGTTCCGAAAAACACCGAGGAAAACAAAATGATGATCCGT
CTGGCGGGCAACAGCGTGGTTCCGGATCAGATTCGTTTCGTGTTTAAGAAACTGTATAGC
GGTTTTAGCAGCATTGTTGATACCACCAGCAACGACGTGGAAATCATTCACTGGAACCCG
CTGCTGGAGAACAACGGCTTCGTTGGTAACATCCACCAGGTGAAGAAACAACGTATTAGC
AACGGCTTTTGCATCGACGGTCGTATTCACGAGAAGCACGTGGTTATCACCCACCGTAAC
CCGATTAACAAAACCCTGTACCCGAACCCGCTGCCGGATAACCACAAGGTTAAAGACCTG
AAGAACGTGGTTACCAACAACGTGACCAAACGTTTCTGGAGCACCCCGTGCCACACCGAT
TTTCGTCGTACCAGCCCGGTGGTTCTGACCAAGCGTCTGCTGACCAACCTGCCGGCGCAA
GTGCGTTATGTTGAAGACAGCGTTCCGGGTTGGCGTCTGAGCGGCGAGTGGTGCCTGTGG
CTGATGGGCTACCACAAAAGCTATGTGACCGGTAGCGAATTC

>YP_001497877
GGATCCATGAGCCTGAAAGCGCTGGAGCTGTTCGCGGGTATCGGTGGCATTACCTACGGC
CTGCGTGGTTATGTGGAACCGATCGCGTTCTGCGAGTACGAAAAAGATGCGGCGGCGTTT
CTGAGCCAGCGTGGCCTGCCGGTTCACGGTGACATTACCAAGTTTGATGCGAGCGTGTAT
AAGACCAAAATCGACATTGTTACCGCGGGTTGGCCGTGCACCGGTTTCAGCACCGCGGGC
AAGGGCACCGGTTTTGATCACGAGGCGAGCGGTCTGTGGACCGAAGTGATCCGTGTGGTT
AAGGAGAGCGAACCGAAATACGTGTTCCTGGAGAACAGCCACGTTCTGGCGCAAACCAAG
AACCTGAAAGTTATCATTCACGACCTGGATGGCCTGGAATACGACACCCGTTGGTGGACC
TGCCGTAGCAACGATGTGAACGTTGGCGCGCACCACAACCGTTATCGTTGGTTCATGCTG
GCGGAGAAGAAAGGTAGCGTGACCAAGTTCGATAAGATCCAGGTTGAAAAGTTCATCTGG
AGCGGCGACTTTAAGGATAAACAAGTGAGCGAGAACAGCCACGAAAACAAGCAACTGATC
AAATTCATGGGTAACAGCGTGGTTCCGGACCAAGTGCGTTACGCGTTTGAGAGCATGAGC
GATATGAAACTGGAAGGCACCCTGGTTAACGACAAGGATGAGATTGTGAAAGTTGGCTAC
AGCAAGGACGGTATCATGTATAAGATTCACGTGGATCACAAGATCATTCCGAAACTGAAC
ATTGTTCTGACCCCGCGTGACCCGCCGGAGGGTCACAAAGCGCGTGAAGATGCGATCATT
AAGAGCCCGATTCTGATGACCTACTGGAACACCCCGGCGTTTTGCTATCACAAAAGCGCG
CGTGGCAGCAAGCTGCTGACCAAGCGTCAGAAAAACAACCTGCACACCCAAATCAAGTTC
TGCCCGGGTGGCAGCGACGATGGCTACCTGAGCGGTCGTTTTTGCGCGTGGCTGATGGGC
TACGGTGACGAATATCTGGGTAACCTGATTGATTATGAATTC

>YP_001497893
GGATCCATGGTACTAAGGGCTTTAGATTTGTTTTCAGGTATCGGCGGTATTACCCACGGA
CTGCGTGAGATTGTGGAGCCGATTGCATTCGTGGAGAAAAACGATGAGGCGCGTAGCTTT
CTGAAGAAGAAACATCCGGAAATCCCGGTTTTTGACGATGTTTGTTCTTTCGACGCGACT
AAGTGGACCCATGTTGATATCATTCTGGCTGGCTGGCCGTGTACCGGTTTCTCTAATGCT
GGCACCAAAACCGGTTTCTCCCACGAAGCGAGCGGTTTGTTCACCGAAGTTGTGCGCATA
ACCGAAGAGTGCCGTCCGAAGTACGTTTTTCTGGAAAACAGCCACACCCTGAGCTTGTTC
GAGAACATTTCCGTGATCGTGAATGCCTTTGATGAACTGGGTTACGACTGTCGTTGGATT
ACGTGCAGAGCGACGTGCGTGGGCGCACTGCACCAGCGTCATCGTTGGTTTTGCCTGGTC
GTCCGCCGTGATATTGAGCTGGACATCGGTATCCCGTATGTGGAGCGCTTTGACTGGATG
ACCAACGAACCGCCTCGCCAGATTCAAAAAAGCAATAAGGAAAACATCCTGCATATTAGC
TTAGCCGGCAACGCGGTTGTTCCGGATCAGGTTCGCTATGCTATGGATATCCTTTGCTCC
TTTGACACCAACAAAGAGTTCTCCTTGTCGAGCCGTGCGAAAAATGGCTTCAGTAAGAAC
GGCGTTATTTTTACCTATGACATTGAGCACCCGGCGCGTGAGCCGCTGAATATCGTGTTG
ACGCCACGTGAGAACGAGACCAGCTTCGTCATCTCTTGTGATCCGAAAAAGGTGCTCACG
AAGTCGGTCATCCAAAAGTTTTGGCCAACGCCGCTGCACAGCTGCAAATACGCTACCAAG
TGCCCGAGAACTCTTACCACCCGTGTGAGCAAGATGCTGTCTGCGTGCGTTGGTTTCAGC
GAAGGTGGCGATAAAAACGGCTACCTGTCCGCGAAATGGGTTCTGTGGCTGATGGGTTAC
GACGACTCATACATGAGCACCGCAGAAAGCAAGTATGAAATGAATCGCGCCTTCTTTCTG
GACATCATCGTAGGTGAACGTGCGAGCGTGTTCGAATTC

>YP_001497965
GGATCCATGACATTAAAAGCTCTAGAATTGTTTGCAGGCATTGCCGGGATCACGCATGGT
TTACGTGGCTTCGTGGAACCGGTTGCATTTGTGGAAATCAACAAAGACGCTCAAGAGTTC
CTATCGACCAAGTTTCCGGACAAGCCGGTATTTGACGATGTTACCAAGTTCTCCAAACGT
GATTTTGATGAGCCGATCGATATGATTACGGGCGGTTTCCCGTGCACCGGTTTCTCTATC
GCGGGTAAGCGTAATGGCTTCGAGCACGCGGAAAGCGGTCTGTTTGGTGAAGTCGTGCGC
ATCACCAAAGAATACATGCCGAAAATGGTTTTCTTGGAGAACAGCGGCATGCTGAGCCAC
AAATATAACCTGGATATTGTCATCAGAAGCATGGATTCCTTGGGCTACGACTGTCGTTGG
GTTACTCTGCGTGCAACCGTGGTAGGTGCTCTGCATACCCGTCATCGTTGGTTTTGCCTG
TGCACCCGCAAAGACCACATTCGCGAGACGCTGATTTGTGATCGTGAGGTGACGAAGTTC
GACTGGGAAAACGATAGACCTCCGATTCAAGTTGATTCACGCTCCTATGAGAACAGCCGT
CTCGTGCGCTTTGCTGGCTACAGCGTCGTTCCGGACCAGATTCGTTATGCGTTCACCGGC
CTGTATACTGGTAACTTTAGCCCGAGCTTTTCCAAGACCTTGGTTCCGGGTAGCCTGGAG
GGTTCCATCTGCTTTAACGAAGACAAGATCACCAATGGTTACTACAAAGACGGTGTTTAT
TACGAGTTCGGCCGTACCGAAACCCACCGTGAGCCGGTGAATATTTTGTTGACGCCACGT
GAGATCCCGAATAAACACAACGGCAAGAAACTGTTGACCCTTCCGGTGACCAAGCGCTTC
TGGTGTACCCCGTGCGCCAGCTATGGTAAGGGTACAGCGGGAGGCCGTGTTCTGACCGAT
CGCTCTAGCCACAGCCTGCCAACGCAGGTGAAATTCTCTCCGGAAGGCGAGGACGGCAAG
CATCTGTCTGGTAAGTTCTGCGCGTGGCTGATGGGTTACGACAAAGAGTATCTGGGAAAT
CTGCTGGAGTACGAATTC

>YP_001498157
GGATCCATGTACTGCATGCTGCGTAGCCTGGACCTGTTCAGCGGCATCGGTGGCAACAGC
TACGCGCTGCGTGATATCCTGAAGCCGATTGCGTATGTGGAGCGTGAACAGCACCTGCGT
GACTTCCTGGGTCGTAAATTTCCGGATGTGCCGATCTTCGACGATGTGGTTACCTTTGAC
ACCCGTAGCGTTAAGGACATTGATATCATTACCGCGGGCTTCCCGTGCACCGGTTTTAGC
ACCGCGGGTAAAGGCGATGGTTTCGAGCACGAAGCGAGCGGCCTGTTTACCGAGGTGGTT
CGTATCGCGAAGGAGCTGGAACCGCGTTTCGTGTTTCTGGAGAACAGCCACACCGTTGCG
CGTATTGAAAACCTGCACGTGATCATTGACGCGTTTGATGTTCTGGGCTACGACTGCCGT
TGGACCACCACCCATGCGACCGCGGTGGGTGCGCCGCAGCAACGTCACCGTTGGTTCTGC
CTGGCGGTGCGTCGTGGTGAAAGCACCGATATCAACATTCCGGACGTTGATCACTTTGAC
TGGACCAACGATGAGCCGGCGAAGCAGATCGAAAACATTGACAGCCGTGGCCGTAAAATC
ATTCAGGCGCTGGGTAACAGCATCGTTCCGGATCAACTGCGTAGCGCGTTCAAGACCATT
ATGAGCATGGACTTTGATGGCAACGAGACCCAAGGTAGCAAGCGTATCCCGCACGGCTAT
AGCATCCGTGGCAAGATCTTCAAGAAAAACATCATCAACACCGAGCGTGAACCGATGAAC
ATCCTGATTTGCCAGAGCGAACCGCCGCAAAAGCACAAAGGCCGTCTGCCGCACATCGAC
AAGAAAATTAAACGTTTTTGGGCGACCCCGGTTAAGGGTACCGCGACCAAGGGTCAGACC
ATCCTGACCCGTCGTAGCCTGCAATGCCTGGGTACCCTGGTGCGTTTCAGCCCGGACGGC
ATTAGCGGTTGGCACCTGAACCCGTTTTGGGTTGGCTACCTGATGGGTTATCGTCGTGAT
TTCTTTGAGGAGTTCGAATTC

>YP_001498439
GGATCCATGTTAGATACACTAGAATTGTTTGCTGGAATCGGCGGCATTACCTATGGTCTG
CGTGGTTTCGCGAAGCCGGCCGCATTTGTTGAGTGGAACGAAGAAGCGAAGAACGTGCTT
AAGCGCCACAAAGTTCCGATCTTTGATGATGTTACTACGTTTGACGCTACCCCGTTTAAA
GACAAAGTTGACATGGTTTCTGCAGGCTGGCCGTGTACCGGTTTCAGCACCGCGGGTCAC
GGCACCGGCTTCTCCCATGAGGCCAGCGGCCTGTTTGTGGAGGTTGTCCGTGTTATCAAA
GAATGCCAGCCGAATTTTGTTTTTCTGGAAAACAGCCACGTTCTCGCGCAAACCCGTTTC
CTAAAGGTGGTGCTGAGCAGTCTGGATGAACTGGGTTACGACGCGCGTTGGATGAGCTGT
AAATCAACCTGTGTTGGCGCAATTCATGAGCGCCATCGCTGGTTTTGCCTGGCTACCAAA
AGAGGTTTTGTGCCACCAACATTGACGAGCCAGGTCAAGCACGAACGTTTCGACTGGGAT
TCGGATGAACCGGCTAAGCAAGTTCCGAAAAACACCGAGGAGAACAAGATGATGATCCGT
CTGGCGGGCAACTCCGTGGTGCCAGACCAGATTCGTTTCGTGTTCAAAAAACTGTACAGC
GGTTTCTCTTCGACCGTTGACACCACTTCCAATGATGTAGAGATCATCCATTGGAATCCG
CTGCTGGAAAACAACGGTTTCGTGGGTAATATTCACCAGGTCAAGAAACAGCGTATTAGC
AATGGCTTCTGCATTGACGGCCGTATCCACGAGAAACACGTGGTCATCACGCATCGTAAT
CCGATCAACAAAACCTTGTACCCGAACCCGCTGCCGGATAATCACAAAGTGAAGGACTTG
AAGAACGTTGTAACCAATAACGTGACGAAGAGATTCTGGTCCACTCCGTGCCATACCGAT
TTTCGCCGTACCTCTCCGGTTGTCTTAACTAAGCGCTTGTTGACGAACCTGCCTGCGCAA
GTACGCTTCGTCGAGGACTCCGTGCCGGGTTGGCGTTTGTCTGGTGAATGGTGCCTGTGG
CTGATGGGTTATAACAAGAGCTATGTTACCGGTAGCGAATTC

>YP_001498713
GGATCCATGGTACTAAGGGCTTTAGATTTGTTTTCAGGCATCGGCGGCATTACCCATGGT
CTGCGTGAAATTGTTGAGCCGATTGCATTTGTGGAAAAAAATGATGAGGCTCGTAGCTTC
CTGAAGAAGAAGCACCCGGAAATTCCGGTTTTCGACGATGTCTGTTCGTTCGACGCCACC
AAGTGGATTGATAAGGTGGATATCATTTTGGCTGGCTGGCCTTGTACCGGTTTCTCTAAT
GCTGGCACCAAGACTGGGTTCTCCCACGAAGCGAGCGGTTTGTTCACCGAGGTCATCCGC
ATGGTCAAAGAGTGCAGACCAAAGTACGTTTTTCTGGAAAACAGCCATACCCTTTCCTTG
TTTGAAAACGTGAACGTAGTGGTGAAAGCGTTTGACGAGCTGGGTTACGACTGTCGTTGG
ATCACCTGCCGCGCGACGTGCGTCGGCGCACTGCACCAACGTCACCGCTGGTTTTGTCTG
GTTGTGCGTCGTGATATCGAACTCGACATCGGCATCCCGTACGTGGAACGTTTTGACTGG
ATGACCAACGAGCCACCGCGCCAGATTCAAAAATCGAACAAAGAGAACATTTTGCATATT
TCCTTGGCGGGTAATGCCGTTGTTCCGGATCAGGTTCGTTATGCTATGGATATCTTGTGC
AGCTTTGACACCAATAAGGAGTTCTCTTTATCCTCTCGTGCGAAAAACGGCTTCAGCAAG
AACGGTGTTATCTTCACCTATGACATCGAGCACCCGGCGCGTGAGCCGCTGAATATTGTG
CTTACGCCGCGTGAGAACGAAACCAGCTTCGTCATCAGTTGCGATCCGAAAAAAGTTCTG
ACCAAGTCCGTGATCCAGAAATTTTGGCCGACGCCGCTGCACAGCTGCAAGTACGCAACC
AAGTGCCCGCGCACTCTGACGACCCGTGTTAGCAAGATGCTGTCAGCGTGCGTGGGCTTT
AGCGAAGGTGGTGATAAAAACGGTTACCTGAGCGCGAAATGGGTTCTGTGGCTGATGGGT
TATGACTATAAATACACACATCATCGCAGTGAATTC

>YP_001498777
GGATCCATGACATTAAAAGCTCTAGAATTGTTTGCAGGCATCGCCGGCATCACGCATGGT
TTGCGCGGTTTTGTAGAACCGATGGCGTTTGTCGAAATTAACAAGGACGCTCAGGAGTTT
CTGTCGACCAAGTTCCCGGACAAGCCGGTGTTCGACGACGTGACCAAATTCAGTAAGCGC
GATTTTGATGAACCGATTGATATGATTACGGGTGGTTTCCCGTGCACTGGCTTCTCCATC
GCGGGTAAACGTAATGGCTTCGAACATGCGGAAAGCGGTCTGTTTGGTGAAGTGGTTCGT
ATTACGAAAGAGTACATGCCGAAAATGGTGTTTCTGGAGAACAGCGGTATGCTGTCTCAC
AAGTATAACCTGGATATTGTTATTCGCAGCATGGATAGCTTGGGCTACGACTGTCGTTGG
GTTACGCTGCGTGCAACCGTTGTTGGTGCGCTGCACACCCGTCATAGATGGTTTTGCCTG
TGTACCCGTAAAGATCATATTAGAGAAACCCTGATTTGCGATCGTGAGGTGACGAAGTTC
GACTGGGAAAACGGCCGCCCTCCGATCCAAGTTGATAGCCGTTCCTACGAGAACTCTCGT
CTGGTGCGTTTCGCCGGTTACAGCGTTGTGCCGGATCAAATCCGTTATGCATTTACCGGC
CTTTATACCGGCAACTTCTCGCCGAGCTTTTCCAAAACCTTGGTACCGGGTAGCTTGGAG
GGCAGCATCTGCTTTAATGAAGACAAGATCACCAATGGTTACTATAAAGACGACGTTTAT
TACGAGTTCGTGCGCACCGAAACCCACCGCGAGCCGGTCAATATCCTGTTGACTCCGCGT
GAAATCCCAAATAAACACAACGGCAAGAAACTGCTGACGCTTCCAGTTACCAAGCGCTTC
TGGTGTACCCCGTGCGCTAGCTACGGCAAGGGCACCGCGGGCGGTCGTGTTCTGACTGAC
CGTTCTAGCCACAGCCTGCCGACCCAGGTCAAATTCTCTCCGGAGGGTGAGGATGGTAAG
CACTTGTCCGGTAAGTTCTGCGCGTGGCTGATGGGTTATGACAAAGAATATCTGGGCAAC
CTGTTAGAGTACGAATTC

>YP_009325790
GGATCCATGGTGCTGCGTGCGCTGGACCTGTTCAGCGGTATCGGTGGCATTACCCACGGC
CTGCGTGAAATCGTTAAACCGGTGGCGTTCGTTGAGAAGAACGACGATGCGCGTGCGTTT
CTGAAGAAAAAGCACCCGACCGTTCCGGTGTTCGACGATGTTTGCAGCTTTGATACCACC
ACCTGGGTGGACAAAGTTGATATCATTACCGCGGGTTGGCCGTGCACCGGCTTTAGCAAC
GCGGGTACCAAGACCGGCTTCAGCCACGAGGCGAGCGGTCTGTTTACCGAAGTGGTTCGT
ATTACCAAAGAGTGCCGTCCGAAGTACCTGTTCCTGGAAAACAGCCACACCCTGAGCCTG
CCGGAGAACGTGAGCGTGGTTGTGAGCGCGTTTGACGAACTGGGCTATGATTGCCGTTGG
ACCACCTGCCGTGCGACCTGCGTGAGCGCGCTGCACCAGCGTCACCGTTGGTTCTGCCTG
GTTGTGCGTCGTGACGTTGAAATCGATTTCGTTGTGAGCTACATTGAAAAATTTGATTGG
AGCGCGAACGAGCCGCCGCGTCAGATCAAAAAGAGCACCAAGGAGAACATCCTGCACATT
AGCCTGGCGGGTAACGCGGTTGTGCCGGACCAAGTTCGTTACGCGATGGATATTCTGTGC
AGCTTCGAGCCGGAAAAATATCCGTGCAGCAGCCGTACCAAAAGCGGTTTCAGCAAGAAC
GGCGTTGTGTTTATGCGTGATGTGAAGCACCCGGCGCGTGAACCGCTGAACATCGTTCTG
CGTCCGCGTGACAACGAGGCGAGCTTTGTTGTGCTGTGCGATCCGAAAAAGGCGCTGACC
AAACCGGTGATTCAAAAGTACTGGCCGACCCCGCTGCACAGCAACAAATATGCGACCAAG
TGCCCGCGTACCCTGACCAAACGTGTGAGCAAGATGCTGAGCGCGTGCGTTGGTTTTAGC
GAGGGTGGCGACAAAAACGGCTATCTGAGCGCGAAGTGGGTTCTGTGGCTGATGGGCTAC
GACGATAAATATATGAGCCCGCTGACCACCATCATTGAATTC

>YP_009665241
GGATCCATGATGCGTAGCCTGGACCTGTTCAGCGGCATCGGTGGCAACAGCTACGCGCTG
CGTGATATTCTGAAACCGGTGGCGTATGTTGAGCGTGAAAAGCACCTGCGTGACTTCCTG
AGCCGTAAATTTCCGGATGTGCCGGTTTTCGACGATGTGGTTACCTTTGACACCAAGAGC
GTGGAGGACATCGATATCATTACCGCGGGCTTCCCGTGCACCGGTTTTAGCACCGCGGGT
AAAGGTGCTGGTTTTGAGCATGAAGCGAGCGGTCTGTTTACCGAGGTGGTTCGTATTACC
AAAGAACTGGTGCCGCGTTTCGTTTTTCTGGAGAACAGCCACACCGTGGCGCGTGTTGAA
AACCTGCACATCATTATCGACGCGTTCGATGTGCTGGGCTACGACTGCCGTTGGACCACC
ACCCATGCGACCGCGGTTGGTGCGCCGCAGCAACGTCACCGTTGGTTCTGCCTGGCGGTG
CGTCGTGAGGAAAGCACCGACATCGCGATTCCGGATGTTGAATATTTTGACTGGACCAAA
GATGAGCCGGTGAAGCAGGTTGAAAACATCGATACCCGTGGCAAGAAAATTATCCAGGCG
ATCGGCAACGGTATTGTGCCGGACCAACTGCGTGCGGCGTTCAAAACCACCATGACCATG
AAGCTGGATGGCATCGAGCTGCGTGGTAACCGTCGTATTAGCCACGGCTACAGCATCCGT
GGCAAGATCTTCAAGAAAAACATCATCATCACCGAGCGTGAACCGATGAACATCCTGATT
TGCCAGAGCGAACCGCCGCAAAAGCACAAAGGTCGTCTGCCGCACATCAAGAACAAAATT
AAGCGTTTCTGGGCGACCCCGGTTAAAGGCACCGCGACCAAGGGTCAGACCATCCTGACC
CGTCGTAGCCTGCAATGCCTGGGTACCCTGGTGCGTTTCAGCCCGGATGGCATCAGCGGT
TGGCACCTGAACCCGTTTTGGGTTGCGTACATCATGGGTTACAAGCAAGACTTCTTTGAT
GAGTTTGAATTC

>YP_009665369
GGATCCATGTTAAATACACTAGAATTGTTTGCTGGAATTGGCGGTATTACCTATGGCCTG
CGTGGTTTTGCGAAACCGGTCGCATTTGTTGAGTGGAACGATGAGGCCAAGAACGTTTTG
AAGCGCCACAAAGTGCCGATTTTTGATGACGTTACGACCTTTGACGCTACCTCTTTTAAA
GACAAAGTCGATATGGTTAGCGCAGGTTGGCCTTGCACCGGTTTCTCCACGGCTGGTCAT
GGCACCGGCTTCAGCCACGAAGCGAGCGGCCTGTTTGTAGAGGTTGTTCGTGTCATCAAA
GAGTGCCAGCCGAATTTTGTGTTCCTGGAAAACTCCCATGTTCTTGCGCAAACCCGTTTC
CTGAAGGTTGTTTTGTCCAGCCTGGATGAACTGGGTTACGACGCCCGTTGGATGAGCTGT
AAATCGACCTGTGTTGGGGCGATTCACGAGCGTCACCGCTGGTTTTGCCTGGCAGTTAAA
CGCGGCTTCGTCCCGCCGACACTGACTTCTCAGGTAAAGCACAAGCGCTTCGACTGGGAT
AGCGATGAACCAGCGAAACAGGTTCCGAAAAATACCGAAGAGAACAAAATGATGATCCGC
TTGGCGGGTAACAGCGTTGTGCCGGACCAGATCCGATTCGTGTTCAAAAAGCTGTACAGC
GGTTTCAGCAGCACCGTTGACACCACTTCTAATGACGTGGAGATCATCCATTGGAATCCA
TTGCTGGAAAACAACGGCTTCGTGGGCAACATCCACCAAGTGAAGAAGCAACGTGTTAGC
AATGGTTTCTGCATTGATGGTCGTATTCACGAAAAACACGTAGTCATCACCCATCGTAAT
CCGATCAACAAGACGCTGTATCCGAACCCGCTGCCGGATAATCACAAGGTGAAGGACCTG
AAGAACGTGGTGACGAACAACGTGACGAAGCGCTTCTGGAGCACCCCGTGCCATACCGAT
TTTCGTCGTACCTCCCCGGTTGTCTTGACCAAACGTCTGCTCACCAACTTACCGGCTCAG
GTGCGTTTTGTGGAAGACTCTGTTCCAGGTTGGAGATTATCCGGCGAGTGGTGTCTGTGG
CTCATGGGTTATCATAAAAGTTACGTAACTGGCAGCGAATTC

>YP_009665506
GGATCCATGGTTCTGCGTGCGCTGGACCTGTTTAGCGGTATCGGTGGCATTACCCACGGC
CTGCGTGAGATCGTTGAACCGATTGCGTTCGTGGAGAAAAACGACGAAGCGCGTAGCTTT
CTGAAGAAAAAGTATCCGGAAATTCCGGTTTTCGACGATGTGTGCAGCTTTGATGCGACC
AAATGGATCGACAAGCTGGATATCATTCTGGCGGGTTGGCCGTGCACCGGTTTCAGCAAC
GCGGGTACCAAAACCGGCTTCAGCCACGAGGCGAGCGGTCTGTTTACCGAAGTTATCCGT
ATTGTGAAAGAGTGCCGTCCGAAGTATGTTTTCCTGGAAAACAGCCACACCCTGAGCCTG
TTTGAGAACGTGAACGTGGTTGTGAAGGCGTTCGACGAGCTGGGTTACGATTGCCGTTGG
ATTACCTGCCGTGCGACCTGCGTTGGTGCGCTGCACCAGCGTCACCGTTGGTTTTGCCTG
GTTGTGCGTCGTGACATTGAACTGGATATCGGTATTCCGTACGTGGAACGTTTCGACTGG
ATGACCAACGAGCCGCCGCGTCAGATCCAGAAGAGCAACAAGGAGAACATCCTGCACATT
AGCCTGGCGGGTAACGCGGTTGTGCCGGATCAAGTTCGTTATGCGATGGACATCCTGTGC
AGCTTCGATACCAACAAGGAATTTAGCCTGAGCAGCCGTGCGAAAAACGGTTTCAGCAAG
AACGGCGTGATCTTTACCTACGACATTGAGCACCCGGCGCGTGAACCGCTGAACATCGTT
CTGACCCCGCGTGAGAACGAAACCAGCTTTGTGATTAGCTGCGATCCGAAAAAGGTTCTG
ACCAAAAGCGTGATCCAGAAGTTCTGGCCGACCCCGCTGCACAGCTGCAAATATGCGACC
AAGTGCCCGCGTACCCTGACCACCCGTGTTAGCAAAATGCTGAGCGCGTGCGTGGGTTTC
AGCGAGGGTGGCGACAAAAACGGCTACCTGAGCGCGAAGTGGGTGCTGTGGCTGATGGGC
TACGATTATAAGTACATGACCAGCGAATTC

>YP_009701829
GGATCCATGCTACACGCTATAGATTTATTTTCAGGAATCGGCGGTATTACCCATGGTTTG
CGTGGTATTGTGGAACCTATTGCGTATGTAGAAAAAAATGATGACGCCAGAGGCTTCCTC
GCGCGTAAACATCCGAATGTTCCGGTGTTTGACGACGTTTGCACCTTTGACGCCACCCCG
TATTTGGGCAAAGTTGATATTATCACTGGCGGCTGGCCGTGCACGGGTTTCAGCACCGCA
GGCAAGGGAACCGGTTTCGAGCACGAGGCCTCCGGTTTATTCACCGAAGTGGTGCGCATC
ACGAAAGAGTGCCAGCCGAAATACTTGTTCTTGGAGAACTCTCACACCCTGGCGGCGTAC
GAGAACATCAACGTTATCGTGAAGGCGTTTGACGACATGGGCTACGATTGCCGTTGGACC
AGCTGTCGTGCTACGTGCGTGGGTGCTCCGCATCAACGTTACCGCTGGTTTTGTCTGGTT
GTGAAAAAAGGCGCTGGCATCGACTTTGAGATCCCGGTCATCGATAAATTCGACTGGGAA
AACAACGAGCCACCGCGCCAGATTGAAAAAAACAACAAGACCAATAAGCTGCGTATCGGT
TTTATGGGCAATGCGGTTGTTCCGGATCAAGTTCGCTACGCAATGACCCTTCTGTCGACG
CTTGAGAACAAGGTTCTGGGTCCGAGCAACACCGATGGTTACAGCATTGACGGTCGTATC
TATACCTTCGTGGTAAAACACCCAACCCGTAAACCGCTGAATATTGTGCTGACGCCGCGT
GAAAACGAAGCGAGCTTTGCGAAAATCTGCGATCCGAAGAAGGTCCTGACCAAGCCAGTC
ACCAAGAAGTATTGGGCAACTCCGGTGTATAATTGTATGAACAGCGCGAAGTGCCCGCGT
ACCCTGACCAAGCGCGTCAGCAACATGCTGTCGGCATGTGTTGGGTTCTCTGAAGGTGGC
AACAAAAACTGGTATATGAACGCCGATTGGATTATGTGGCTGATGGGTTACGAGCCGGGT
TACCTGAGCCACGAATTC

>YP_009701874
GGATCCATGACCTACCGTACCCTGGAGCTGTTTGCGGGCGTTGGTGGCATCACCCATGGT
CTGCGTGGCATTAGCACCCCGGTTGCGTTCGTGGAGATCAACAAGGACCCGCAGCAATTT
CTGAAGACCAAATTCCCGGAAGCGAGCGTTTTTGACGATGTGACCAAGTTTACCAAAGAG
GACTTCCCGGAAACCATCGATATGATTACCGCGGGTTTTCCGTGCACCGGCTTCAGCATC
GCGGGTAGCCGTACCGGCTTTGAGCACAAGGAAAGCGGTCTGTTCGCGGATGTGGTTCGT
ATTACCGAGGAATATAAGCCGAAACTGGTTTTCCTGGAAAACAGCCACATGCTGAGCCAC
ACCTACAACCTGGACGTGGTTGTGAAGACCATGGATAAGATCGGTTACTTCTGCAAGTGG
ATCACCTGCCGTGCGAGCGTTGTTGGTGCGCACCACCAGCGTCACCGTTGGTTCTGCCTG
GCGACCCGTAAGGACTACGTGCCGGAGAAAATCGAAGTTAGCGTGAACGCGACCAAGTTC
GATTGGGAGAACAACGAACCGCCGTGCCAGGTTGAGAACAAAAGCTATGAAAACAGCACC
CTGGTGCGTCTGGCGGGTTACAGCGTTGTGCCGGACCAAATTCGTTATGCGTTCACCGGT
CTGTTTACCGGCGATTTCGAGAGCAGCTGGAACACCACCCTGACCCCGGGTACCATCACC
GGTACCGAACACGTTAAGATGAGCGGCAACTACGACAAAGTGATTAACGGTTATTGCGTT
GATGGCGTGTACTATGAGTTTAGCCGTAAGGAAACCCACCGTGCGCCGCTGAACATCAGC
GTTAAACCGCGTGCGATTCCGGAGAAGCACAACGGTAAAACCCTGGTGGACCGTGAAATG
ATCAAGAAATACTGGTGCACCCCGTGCGCGAGCTATGGTACCGCGACCGCGGGTTGCAAC
GTTCTGACCGATCGTCAGAGCCATGCGCTGCCGACCCAAGTTCGTTTCAGCTATCGTGGT
GTGTGCGGTCGTCACCTGAGCGGTATTTGGTGCGCGTGGCTGATGGGCTACGACCGTAAA
TACCTGGAGTATCTGGTGGATTATAACGAATTC

>YP_009702000
GGATCCATGCTGCGTGCGCTGGACCTGTTCAGCGGCATCGGTGGCATTACCTACGGCCTG
CGTGGTATCGTTACCCCGGTGGCGTATGTTGAGAAGAACAAAGACGCGCGTGAATTTCTG
CAGAAGAAACACCCGGATGTTCCGGTGTTCGACGATGTTTGCGCGTTTGACGCGACCGAG
TGGAAGGGCAAAGTGGATATCATTACCGGTGGCTGGCCGTGCACCGGTTTCAGCATCGCG
GGTAAAGGCGCGGGTTTCGAGCACGAAGCGAGCGGTCTGTTTAGCGAGGTGGTTCGTATC
ACCAACGAGTGCGAACCGCGTTACCTGTTCCTGGAAAACAGCCACATTCTGAGCAAGAAA
ACCAACGTTAGCGTGGTTGTGAACGCGCTGGACAAGCTGGGCTACGATTGCAAATGGCTG
ACCTGCCGTGCGACCAACGTGGGTGCGCCGCACCAACGTCACCGTTGGTTCTGCCTGGTG
ATCAAGCGTGGCGCGAGCGTTATTACCGACCTGCCGCTGGTTGACACCTTTAACTGGAGC
AGCGGCGAGCCGGACAAGCAGATCAAAAACTATACCCCGATTAACCGTACCGTTAGCAGC
TGGCTGGGTAACGCGGTTGTGCCGGACCAAGTGCGTTACGCGTTCACCACCCTGAACACC
ATGAGCACCGAAATCGCGACCTATCAGCACGATATTAACAAGTTCCAAAACTGCGGTTTT
AGCCTGAAAAGCGAGATCTACTGGAGCCGTATCAAGATTCAGGAACGTCCGCCGCTGTAT
ATTACCCTGACCCAAGAGAACCCGCCGAACAAGCACAAAGCGACCACCCCGCACATCAAA
ACCATTACCAAGAAATTCTGGAGCACCCCGGTTTTTAGCCACAACCCGAGCGGCGTGAAC
TTCCTGACCTACCGTAGCAGCAAGCTGCTGGGCACCCAGGTTAAATTCGCGGAAGGTGGC
TTTGCGGGCATGTATGTGAACCCGGATTGGGTTCGTTGGCTGATGGGTTACCCGGAGGAC
TATTTTACCCTGGCGTACAAGGATCACGAAACCCAAAGCGAATTC
