## Supplementary Table S1 for "Identification, expression, and purification of DNA cytosine 5-methyltransferases with short recognition sequences"

**Supplementary Table S1. Summary of the analysis of candicate DNMTs**

| Entry | Vector | Gene | Cloning | Western*1 | E. coli WGBS*2 | Protein purification*3 | In vitro activity*4 | Assay5* |
|---|---|---|---|---|---|---|---|---|
| 1 | pCZS | M.CviPI | + | + | GC | ++ | + | |
| 2 | pCZS | M.CviPII (NDel15) | + | + | CC | ++ | + | HaeIII |
| 3 | pCZS | M.CviQIX (NDel15) | + | + | CC | ++ | + | |
| 4 | pCZS | NP_048873 | + | ++ | TCTG | ++ | + | WGBS |
| 5 | pCZS | NP_048886 | + | ++ | TCG | + | - | NruI |
| 6 | pCZS | NP_049039 | + | ++ | GCY | - | - | |
| 7 | pCZS | YP_001426001 | + | ++ | CG | + | + | HpaII |
| 8 | pCZS | YP_001426380 | + | ++ | CC | ++ | - | HpaII |
| 9 | pCZS | YP_001426521 | + | ++ | CG | + | + | HpaII |
| 10 | pCZS | YP_001497284 | - | - | - | - | - | |
| 11 | pCZS | YP_001497607 | + | ++ | - | - | - | |
| 12 | pCZS | YP_001497877 | - | - | - | - | - | |
| 13 | pCZS | YP_001497893 | - | - | - | - | - | |
| 14 | pCZS | YP_001497965 | - | - | - | - | - | |
| 15 | pCZS | YP_001498157 | - | - | - | - | - | |
| 16 | pCZS | YP_001498439 | - | - | - | - | - | |
| 17 | pCZS | YP_001498713 | + | ++ | TCG | - | - | |
| 18 | pCZS | YP_001498777 | - | - | - | - | - | |
| 19 | pCZS | YP_009325790 | + | ++ | TCG | + | - | HpaII |
| 20 | pCZS | YP_009665241 | + | + | - | - | - | |
| 21 | pCZS | YP_009665369 | - | - | - | - | - | |
| 22 | pCZS | YP_009665506 | + | ++ | TCG | - | - | |
| 23 | pCZS | YP_009701829 | - | - | - | - | - | |
| 24 | pCZS | YP_009701874 | + | ++ | GCY | + | + | HaeIII, HaeII, HhaI |
| 25 | pCZS | YP_009702000 | - | - | - | - | - | |
| 26 | pBZS | M.CviPI | + | ++ | GC | + | + | HaeIII |
| 27 | pBZS | M.CviPII (NDel15) | + | ++ | CC | - | - | HaeIII |
| 28 | pBZS | M.CviQIX (NDel15) | + | ++ | CC | + | + | HaeIII |
| 29 | pBZS | NP_048873 | + | - | - | - | - | |
| 30 | pBZS | NP_048886 | + | - | - | - | - | |
| 31 | pBZS | NP_049039 | + | ++ | - | - | - | |
| 32 | pBZS | YP_001426001 | + | ++ | CG | ++ | + | HpaII |
| 33 | pBZS | YP_001426380 | + | - | - | - | - | |
| 34 | pBZS | YP_001426521 | + | ++ | - | - | - | |
| 35 | pBZS | YP_001497284 | + | ++ | CG | - | - | |
| 36 | pBZS | YP_001497607 | + | + | CNG | ++ | + | WGBS |
| 37 | pBZS | YP_001497877 | + | ++ | CC | - | - | |
| 38 | pBZS | YP_001497893 | + | ++ | - | - | - | |
| 39 | pBZS | YP_001497965 | + | ++ | GC | + | + | HaeIII |
| 40 | pBZS | YP_001498157 | + | ++ | CG | - | - | |
| 41 | pBZS | YP_001498439 | + | ++ | CNG | ++ | + | WGBS |
| 42 | pBZS | YP_001498713 | + | ++ | - | - | - | |
| 43 | pBZS | YP_001498777 | + | ++ | GC | - | - | |
| 44 | pBZS | YP_009325790 | + | ++ | - | - | - | |
| 45 | pBZS | YP_009665241 | + | - | CG | - | - | |
| 46 | pBZS | YP_009665369 | + | ++ | CNG | ++ | + | WGBS |
| 47 | pBZS | YP_009665506 | + | ++ | - | - | - | |
| 48 | pBZS | YP_009701829 | + | + | CG | ++ | + | HpaII |
| 49 | pBZS | YP_009701874 | + | ++ | GCY | - | - | |
| 50 | pBZS | YP_009702000 | + | ++ | RGCA | ++ | + | WGBS |

*1 After induction of the protein expression, bacterial lysate was served for western blotting using anti-strep II tag antibody. ++: stong and shap signal detected, +: signal detected

*2 Target motif identified by WGBS of E. coli genomic DNA extracted from cells expressing the protein

*3 ++: purified protein was almost homogenous, +: protein signal was detected but unpurities were also observed

*4 +: DNA methylation was detected with the methylation-sensitive restriction enzyme or WGBS

*5 Assay types used for the methylation activities. Umnethylated lambda DNA was used for the model.
